# Humans compress order-based structure into short mental programs

**DOI:** 10.64898/2026.05.14.725160

**Authors:** Elyes Tabbane, Santiago Figueira, Lucas Benjamin, Stanislas Dehaene, Fosca Al Roumi

**Affiliations:** Cognitive Neuroimaging Unit, CEA, INSERM, Universite Paris-Saclay, NeuroSpin Center, Paris, France; Paris Brain Institute; Universite Paris-Cite; Departamento de Computacion, Facultad de Ciencias Exactas y Naturales, Universidad de Buenos Aires, Buenos Aires, Argentina; Instituto de Ciencias de la Computacion (ICC), CONICET-Universidad de Buenos Aires, Buenos Aires, Argentina; Cognitive Neuroimaging Unit, CEA, INSERM, Universite Paris-Saclay, NeuroSpin Center, 91191 Gif/Yvette, France; Institut National de la Sante et de la Recherche Medicale, Institut de Neurosciences des Systemes; DEC,Ecole Normale Superieure-Universite Paris; Cognitive Neuroimaging Unit, CEA, INSERM, Universite Paris-Saclay, NeuroSpin Center, 91191 Gif/Yvette, France; College de France, Universite Paris Sciences Lettres (PSL), 11 Place Marcelin Berthelot, 75005 Paris, France; Cognitive Neuroimaging Unit, CNRS ERL 9003, INSERM U992, CEA, Universite Paris-Saclay, NeuroSpin Center, 91190 Gif/Yvette, France

## Abstract

How do humans store sequences that far exceed working memory capacity? Previous work showed that visuo-spatial sequences with geometric regularities and binary auditory sequences are compressed into mental programs made of nested compositions of simple operations provided by the Language of Thought model. Here we ask whether this principle extends to a more abstract domain: order-based structure, such that sequences would be coded by how items repeat and in what order, leveraging ordinal positions in memory. For instance, imagine the sequence of tones CCDDEECCDDEE and the sequence of colors RRGGBBRRGGBB (R: red, G: green and B: blue). They share the same structure, which lets you store 12 items as “3 pairs, repeat”. In two experiments, participants memorized and reproduced 12-item sequences of spatial locations; in a third, they rated the complexity of those same sequences. The order-based Language of Thought description of the sequences predicted recall accuracy with high precision (r = .96), outperforming several alternative sequence complexity measures, but also participants’ own subjective complexity ratings. Beyond accuracy, the inferred programs predicted the structure of motor production of sequences: inter-click intervals slowed down and errors increased at the constituent boundaries specified by the model. Finally, we identified a set of elementary operations (mirroring, interleaving) that are also building blocks of such mental programs. Together these findings indicate that when a sequence affords an order-based structure, it is encoded as a mental program in human memory. We thus extend the validity of the Language of Thought framework to an abstract code, in which structure relies on the order in which items appear rather than modality-specific regularities.

**Author Summary:** Short-term memory holds only a few items at once. Yet we routinely memorize sequences far exceeding this capacity. How is this possible? We posit the brain acts like a programmer: rather than storing each element individually, it compresses sequences into short mental programs, just as a programmer writes “repeat ABC four times” instead of typing ABCABCABCABC. Such programs have been shown to describe domains from geometry to proto-musical sequences. We show that they also operate on a more abstract dimension: the order in which items appear. In two experiments, participants memorized and reproduced sequences of flashed positions on a screen; in a third they rated their complexity. Sequences described by shorter programs were remembered far better, and when reproducing them, participants paused at the boundaries between program parts, revealing the internal mental description. Strikingly, program length predicted memory performance better than participants’ own complexity ratings. Finally, we identified a set of elementary operations (mirroring, interleaving) that are building blocks of such mental programs. Our findings suggest that the brain leverages order-based structure to compress information in memory.

## Introduction

Consider a melody of twelve notes that has the following structure: *AABBCCAABBCC*. Remembering twelve sounds surpasses typical working memory limits [1–3]. Yet humans can easily memorize this sequence as a short mental program “repeat twice: 3 pairs of items”. How are humans bypassing these constraints? It has been proposed that much of our cognition can be understood as a drive toward simplicity by compressing information and eliminating redundancy [4]. Accordingly, human cognition recodes presented information into a more compact, storage-efficient format, a process known as compression. And because compression relies on the recognition of regularities, memory load depends less on the number of items than on the structure linking them [5–9].

Recent work suggests that the human brain encodes structured experiences as generative programs that compress information based on its underlying structure. Building on the Language of Thought (LoT) hypothesis [10], recent accounts [11–19] posit that humans can represent structured experience as mental programs, in which elementary operations, called primitives, are composed to capture the relational structure. The LoT hypothesis assumes that, to represent a sequence, the brain selects the shortest possible description with the given set of available primitives. The length of this shortest program (or minimal description length) is called LoT-complexity. Operationalizing stimuli as generative programs within the LoT framework allows two main analyses: (1) LoT-complexity can be compared to other complexity metrics, to test which better reflects humans’ cognition; (2) programs specify constituent structure, and therefore not only predict a sequence’s difficulty but also its representational structure and the hierarchical grouping of elements. The LoT framework has been applied to a range of domains including numerical concept acquisition [12] and the induction of abstract visual concepts [13]. It has recently emerged as a powerful model for understanding how the human brain encodes sequences, supporting the view that visuo-spatial sequences with geometrical regularity and binary auditory sequences are compressed into hierarchical mental programs [14–19]. LoT-complexity proves to be a good predictor of participants’ behavior [14–19], but also of brain activity [14,15,19]. Notably, evidence shows that mental programs do not recruit regions associated with natural language, but rather a fronto-parietal network that overlaps with the one for mathematical reasoning [15,19]. Furthermore, the primitive operations could be decoded from brain activity as well as their composition into programs’ components [14].

Together these results suggest that human cognition relies on an efficient capacity for representing structured information, coding information as generative mental programs [17]. However, it remains unknown whether similar mental programs can compress sequences solely from their order-based structure, that is, when structure is defined not by physical coordinates (like position in the visual field), but by abstract coordinates coding the order in which items appear and repeat (see **Fig 1A** and **1B**). For instance, consider the following sequences aabbccaabbcc and ffeeddffeedd, where letters from ‘a’ to ‘f’ refer to spatial positions spanning the vertices of a hexagon (**Fig 1B**). While the specific spatial coordinates differ, both sequences share an identical order-based structure: 112233112233. To capture this shared structure, the brain must move beyond a perception-based map of where each item was flashed to a more abstract code in which items relate to a temporal feature [20–24] and are relabeled by their ordinal code. We hypothesize that regardless of the physical identity of the stimuli (e.g. spatial locations, auditory tones, or visual objects) whenever a sequence’s structure lies in the ordinal dimension, a LoT tuned to order-based structure can efficiently describe how participants encode it. Consequently, while we utilized a visuo-spatial experimental setting, the LoT operates completely blindly to physical geometry by only considering items as abstract ordinal memory slots (see **Fig 1A**). Our order-based implementation of LoT shares its architecture with previous work [14,16,19] but now applies over ordinal code rather than physical space. Hereafter, we refer to this implementation as the *order-based LoT model*. In computer-science terms, it corresponds to a domain-specific language (DSL) for order-based structure.

**Figure 1.**
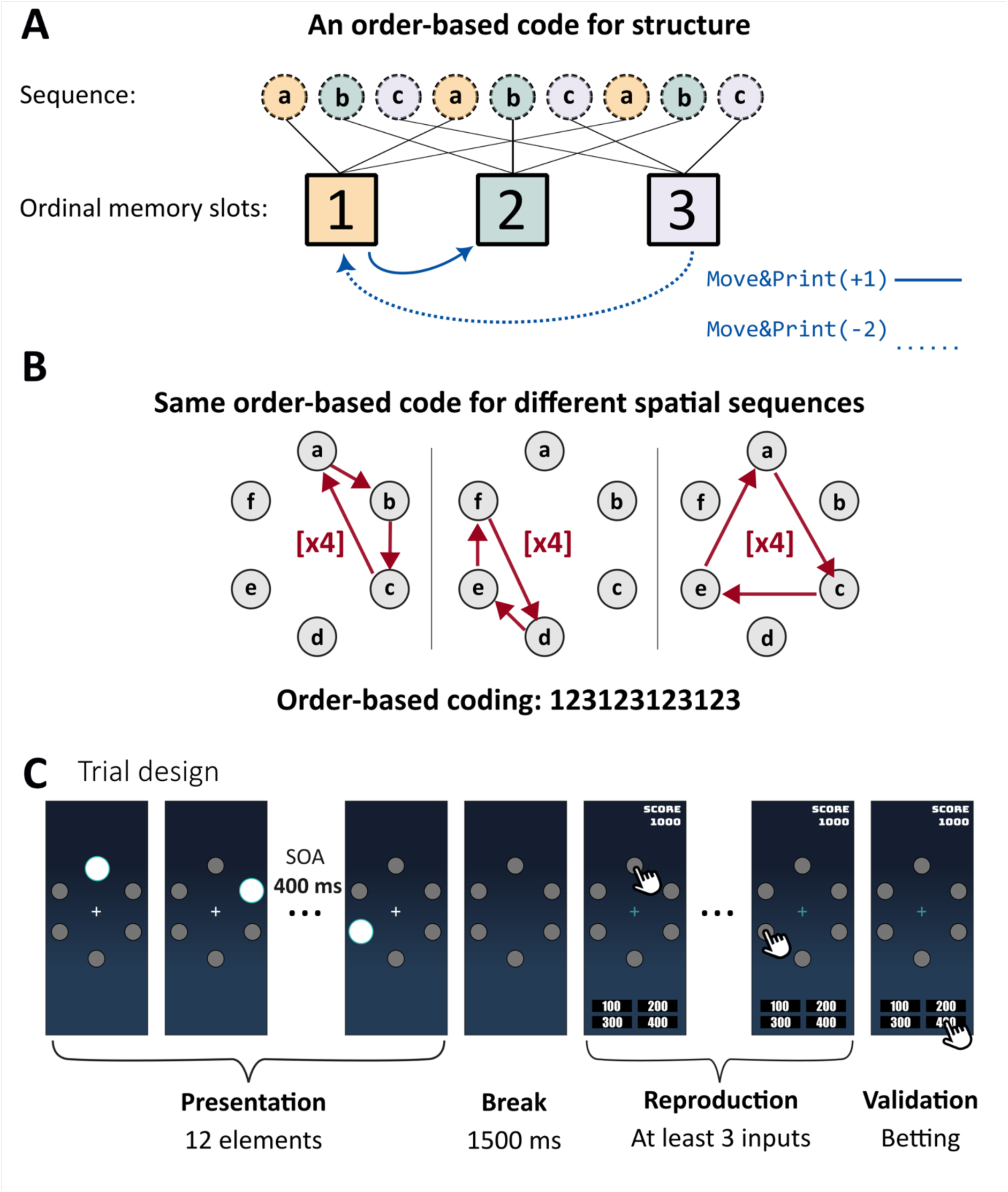
Order-based coding of sequences and experimental design. **(A)** Ordinal recoding. The items of a visuo-spatial sequence are recoded into ordinal memory slots according to their order of first appearance. The primitive Move&Print(n) moves the point of evaluation by n slots and outputs the item stored in the slot it reaches; Move&Print(0) outputs the current item again. **(B)** The same order-based code for different spatial sequences. Three sequences visiting different vertices of the hexagon (abcabcabcabc, fedfedfedfed and eaceaceaceac), each repeating a three-item path four times (×4), share the same order-based code, 123123123123. Spatial positions attributed to 1, 2 and 3 were drawn randomly among the vertices of the hexagon. **(C)** Trial design. Each trial began with the presentation of a 12-item sequence of flashed yellow circles at six fixed spatial positions arranged hexagonally (SOA 400 ms). After a 1500 ms retention interval, the fixation cross turned blue to indicate the start of the response phase, and participants reproduced the sequence by clicking on the placeholders (at least three responses). They validated their sequence by choosing a bet (100–400 points) reflecting their confidence, and feedback was given as points gained or lost. In the subjective complexity rating task (**Experiment 3**), the presentation phase was identical, but participants rated the memorability of each sequence on a 7-point scale (1: Very Simple to 7: Very Complex) and no feedback was provided. The full set of primitives and a step-by-step example of program execution are shown in Fig A in S1 Text.

We predicted that if order-based structure is represented as mental programs, working memory performance should be determined by the order-based LoT-complexity (denoted simply as LoT-complexity hereafter) rather than the number of items in the sequences. To test this prediction, we conducted a series of experiments using a smartphone-based paradigm, requiring participants to memorize and reproduce visuo-spatial sequences of 12 items [25,26]. Critically, we randomized the spatial positions used at every trial such that any geometrical regularity effect would be averaged out (see **Fig 1B**). Across three experiments, we showed that LoT-complexity predicts recall performance better than competing models –including Shannon entropy over transition probabilities, Lempel-Ziv complexity, and chunking models – indicating that memory load is best accounted for by mental programs describing order-based regularities. We further fitted the relative weights of several primitives on one cohort and validated them on another. The obtained model further predicts the fine-grained temporal structure of participants’ production: inter-click intervals slow at constituent boundaries of the programs. By extending the stimulus set to a broader range of order-based regularities, we challenged the scope of the order-based LoT model and identified additional primitives required to account for spontaneous compression of a wide variety of structures. Finally, subjective complexity ratings partially deviated from LoT complexity, indicating that the compression operates without being entirely introspectively accessible. Together, these results support the view that working memory for order-based sequences relies on a program-like representation that shapes both memory load and the timing of motor execution, while only being partially accessible to introspection.

## Experiment 1. Repetitions foster memory compression

### Methods

Our experiments were performed online on a smartphone. Available languages were French and English. The online experiment was built using JavaScript (ES6), HTML (v5) and CSS (v3).

#### Participants recruitment and ethics

Participants’ data were anonymized. The experiment was conducted with ethical approval from the Research Ethics Committee (CER) of the University of Paris-Saclay (reference: 2019-063). Before starting, participants read an information page describing the task, its duration, the data collected (responses and response times only), the possible use of the results in scientific publications, and the absence of risk beyond that of daily life; minors were asked not to take part. Informed consent was obtained electronically: participants consented by starting the experiment, and no written consent was collected, as the data were fully anonymous. Participation was voluntary and unpaid. We recruited 99 participants via social media (X). To ensure data quality in an unmonitored setting, we excluded 4 participants who either exceeded a 10-minute session threshold or produced unshown positions on each trial and 13 participants who took the experiment more than once. The final sample included 82 participants (mean duration = 5 minutes 54 seconds).

#### Sequences tested in Experiment 1

**Experiment 1** focused on 12-item sequences that were structured with repetition, and the corresponding controls. The control sequences used the same elements as the structured ones but were arranged in a different temporal order. Sequences were chosen to sample the main ways in which repetition can structure a 12-item sequence: the repetition of a set of 2, 3 or 4 items (Rep-2, Rep-3, Rep-4), and the repetition of a repetition (Rep-Nested), in which three doubled items form a block that is itself repeated. The control sequences remove this structure at one or both hierarchical levels: Rep-Global keeps the repetition of a whole block but removes the local doubling, Rep-Local keeps the local doubling but removes the repetition of the block, and CRep-2, CRep-3 and CRep-4 remove repetition altogether. Controls were constructed by hand rather than by random shuffling, selecting among reorderings of the same elements the arrangement that maximized LoT-complexity while eliminating group-level repetitions. Here, we present the sequences and their description provided by the order-based LoT model (see section *Parameter search*).

***Rep-2: 121212121212:***

Repetition applied to a set of 2 elements, 6 times.

***CRep-2: 122221121112.***

***Rep-3: 123123123123***:

Repetition applied to a set of 3 elements, 4 times.

***CRep-3: 123132231213:***

Control obtained from shuffling groups of 3 items in the sequence.

***Rep-4: 123412341234:***

Repetition applied to a set of 4 elements, 3 times.

***CRep-4: 123432411423:***

Control obtained from shuffling groups of 4 items in the sequence.

***Rep-Nested: 112233112233:***

3 items, each doubled. The entire block is repeated twice.

***Rep-Global: 123132123132:***

Control for Rep-Nested with no local repetitions. Repetition of 123132, twice.

***Rep-Local: 112233113322:***

Control for Rep-Nested with no global repetition but with local ones (doubling).

Rep-Global and Rep-Local were designed by hand to knock out a single hierarchical level of the Repeat primitive in the Rep-Nested structure.

### Spatial layout

Each pattern could use any subset of the six vertices of the hexagon, which were chosen randomly (see **Fig 1B**). Thus, each pattern had a large number of possible realizations in terms of actual spatial coordinates used. 30 possible realizations for Rep-2 and CRep-2 (6×5); 120 for Rep-3, CRep-3, Rep-Nested, Rep-Global and Rep-Local which all use three spatial positions (6×5x4); 360 for Rep-4 and CRep-4 (6×5x4×3).

#### Experimental design

The following instructions were shown to the participants. “*A sequence of points will be presented to you. Try to memorize it. After a short delay, you’ll be asked to reproduce it. Please use your index finger. Once you are done, push one of the buttons at the bottom of the screen to bet on your guess. The higher your bet, the higher your potential gains or losses! You have to enter at least 3 points before you are able to bet.*”

The experiment started with a few training trials to familiarize participants with the task design. Participants were not explicitly instructed that sequences contained 12 items.

The figure on screen consisted of 6 small white circles (50 pixels radius) positioned hexagonally and a central fixation cross (50 pixels width and height). When a sequence was presented, bigger yellow circles (90 pixels radius) were flashed (300ms) on top of the smaller circles. When participants were clicking on a small circle to reproduce the sequence, a bigger yellow circle was flashed at the same location and a ‘click’ sound was presented to provide visual feedback and inform participants their response was recorded.

##### Trial flow: delayed reproduction with betting (Fig 1C)

###### Presentation phase

The 12 items of the target sequence were presented with SOA 400ms. Stimulus duration on screen was 200ms.

###### Maintenance and response phase

A 1500ms retention interval followed sequence presentation. The fixation cross changed from white to blue to indicate the end of this delay and the start of the response phase. Simultaneously, 4 bet buttons with values 100, 200, 300 and 400 appeared at the bottom of the figure. Once participants had clicked on at least 3 positions, they could validate their response by pressing one of the bet buttons. After 15 positions, further responses were not recorded and the circles became unresponsive.

###### Feedback phase

Following score feedback, a 750ms delay occurred after participants clicked “next” to begin the next sequence.

###### Sequences

To even out geometrical regularities, spatial positions were randomized on each trial. 9 distinct sequences were presented 2 times for a total of 18 trials.

###### Betting and Scoring

Starting with 1,000 points, participants validated each trial by betting 100, 200, 300, or 400 points. A perfect sequence reproduction earned the full bet. If participants’ response was incorrect but similar to the target sequence, participants earned a fraction of the bet. Otherwise, the amount of points they lost scaled with the bet and the dissimilarity of their production relative to the correct sequence, as computed via the edit distance (see below). Token errors (see below) were penalized heavily: they led to the loss of the chosen bet value. Feedback was designed in this manner to encourage participants to try their best even on difficult trials.

###### Feedback messages

The feedback screen presented the number of points gained or lost (- 400 to +400), as well as a short encouraging message. When participants lost points, the feedback message was encouraging (“That was close”, “Almost got it”, etc.). When participants earned points, they received praise (“perfect”, “amazing”, “genius”, etc.) according to their performance.

#### Performance Measures

*A Token Error* was defined as a discrepancy S≠R between the set of unique stimuli presented (S) and the set of unique stimuli recalled (R). This classification encompasses both *intrusions* (producing a position not in S) and *omissions* (not producing all the positions of S).

*Forward span* was defined as the number of correctly recalled items in serial order preceding the first error on a given trial, averaged across trials for each participant (i.e., mean correct prefix length before the first error).

As an edit distance measure, we used the *Damerau-Levenshtein (DL) distance* [27] to quantify sequence dissimilarity. It measures the minimum number of operations: insertion (*AB to ABC)*, deletion (*ABCC to ABC)*, substitution (*ABB to ABC)*, or transposition (*ACB to ABC*), needed to transform one string into another. This provides a finer-grained metric than error rate. For instance, if a participant reproduces the sequence 121212121212 as 212121212121, a sequence shift results in a 100% error rate but an edit distance of only 2. All edit distances were computed with a custom Python implementation of the standard dynamic-programming algorithm for the restricted Damerau-Levenshtein distance (also known as optimal string alignment), in which insertions, deletions, substitutions and transpositions of adjacent items all have a cost of 1 (function dl_distance in our analysis code; see Availability statement).

Retroactive interference was operationalized as error rate on the shared first constituent of each structured sequence and its paired control (e.g., the opening “123…” common to Rep-3 and CRep-3). Because these elements are identical, differences in recall accuracy of these first items reflect a backward influence of the sequence’s global structure on memory for its earliest elements, rather than differences in local encoding difficulty.

#### Statistical tests

##### Paired conditions comparison

All of our compared samples were dependent (same participants but on different conditions). Parametricity of data was assessed before each comparison, using a Shapiro-Wilk test. When p < .05 we ran a Wilcoxon signed-rank test, and when p > .05 we ran a paired T-test.

##### Effect size

For paired T-tests, we report Cohen’s d as a measure of effect size. For non-parametric comparisons between paired conditions (e.g., edit distances, RTs), we used the Wilcoxon signed-rank test. To estimate the magnitude of the effect, we report the effect size r, computed as r = Z / √N where Z is the standardized test statistic and N is the number of non-zero difference pairs. This measure can be interpreted similarly to a Pearson’s correlation coefficient, with r = .1, .3, and .5 corresponding to small, medium, and large effect sizes, respectively [28].

### Results

The 82 participants accurately remembered the spatial positions (98.64% correct) but often failed to reproduce their temporal order (21.34% correct, SEM = 2.12%). Participants showed a strong primacy effect (first three items: +19.63%, +14.62%, +14.82% above mean accuracy) but no recency effect. We assessed participants’ capacity to bind spatial and ordinal information by computing the forward span (see *Methods*) on control sequences. No significant difference was found between CRep-3 (2.21, SEM = 0.21) and CRep-4 (2.09, SEM = 0.19; W = 1053, p = .90), suggesting a baseline capacity of ∼2 items for unstructured sequences. In contrast, forward span was higher and differed between Rep-3 (7.73, SEM = 0.46) and Rep-4 (6.41, SEM = 0.47; W = 814, p = .03), reflecting the benefit of order-based structure on memory capacity.

#### Sequence Performance and Accuracy

We compared performance on structured sequences (Rep-2/3/4) against control sequences (CRep-2/3/4). We excluded Rep-Nested and its controls from this analysis, as the latter still possessed underlying order-based structure. Structured sequences are overall better reproduced than controls (see **Fig 2A**). Order-based structure was a primary determinant of success across both metrics. Structured sequences exhibited significantly lower error rates (66.67%, SEM = 3.11%) than control sequences (97.56%, SEM = 1.23%; W = 0.00, p < .001, r = .88). Edit distance was lower for structured than control sequences (structured sequences edit distance = 1.95, SEM = 0.12, control sequences edit distance = 4.55, SEM = 0.1; r = .87, Wilcoxon signed-rank test p < .001). Every structured sequence was significantly better memorized than its paired control, yielding large effect sizes (r ≥ .85) and high significance (p < .001) for both error rates and edit distance (see **Fig 2A**). A similar pattern emerged for recursive repetitions. Error rates and edit distance showed that the Rep-Nested sequence had a significant memory advantage over Rep-Local (all p < .005; r_error = .45 and r_edit_distance = .59), Rep-Global (all p < .001; r > .80) and CRep-3 (all p < .001; r > .86). Notably, performance on the Rep-Nested sequence did not differ significantly from the Rep-3 sequence in both error rate and edit distance (all p > .05). Detailed results for pairwise comparisons are provided in Tables B–D (error rate) and Tables E–G (edit distance) in S1 Text.

**Figure 2.**
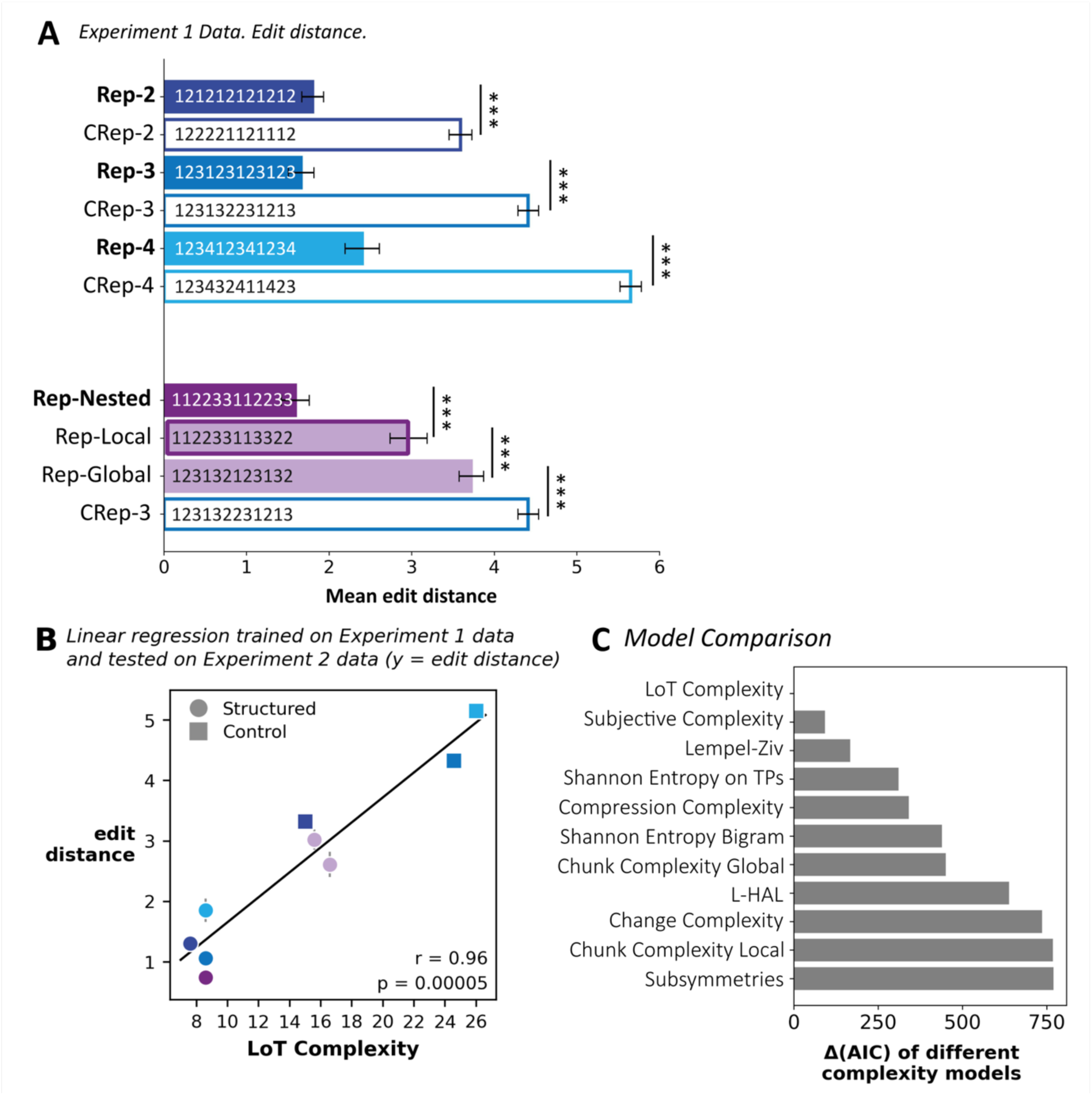
Temporal Language of Thought Supports Sequence Recall. **(A)** Performance, measured as edit distance, reflects temporal structure. Data reported here belong to **Experiment 1**. Error bars indicate SEM of per-participant averages. Asterisks denote statistical significance of per-participant mean edit distance distributions, assessed with a Wilcoxon signed-rank test (*** : p < .001). **(B)** LoT Complexity predicts performance. Model parameters were estimated from **Experiment 1** data and evaluated on **Experiment 2**, which included a larger set of sequences and the full set from **Experiment 1** for replication. The Pearson correlation and the regression line are here computed on the mean edit distance per sequence. Color code corresponds to color on panel A. **(C)** LoT outperforms alternative models of sequence complexity. The panel shows Akaike Information Criterion (AIC) differences between mixed-effects models based on LoT Complexity and those based on competing sequence-complexity metrics.

Participants were significantly faster at producing structured sequences (6147ms, SEM = 141ms) than control sequences (6494ms, SEM = 150ms; W = 1047, p = .003, r = .33), though the advantage of order-based structure decreased as the number of tokens increased. Specifically, while Rep-2 (W = 662, p < .001, r = .53) and Rep-3 (W = 1142, p = .01, r = .29) showed significant speed benefits, the effect was not significant for Rep-4 (W = 1488, p = .32, r = .11). Regarding recursive repetition, Rep-Nested (6232ms, SEM = 147ms) did not statistically differ from Rep-3 and CRep-3 (both p > .05), but it was significantly faster than Rep-Local (6673ms, SEM = 179ms, W = 1002, p = .001, r = .36) and Rep-Global (6638ms, SEM = 190ms, W = 1243, p = .03, r = .23). Detailed pairwise comparisons are provided in Tables H– J in S1 Text.

To test if sequence structure retroactively influences memory, we compared error rates on the identical starting items (e.g., ‘123’) shared by structured and control sequences. All tested pairs differed significantly (p < .001). Detailed results for pairwise comparisons are provided in Table M in S1 Text. The largest differences appeared in Rep-3 and Rep-4. For Rep-3, the ‘123’ constituent error rate was 26.22%, SEM = 3.8% (structured) vs. 57.93%, SEM = 3.93% (control; W = 312, r = .64). For Rep-4, the ‘1234’ constituent error rate was 40.24%, SEM = 4.46% (structured) vs. 66.46%, SEM = 3.98% (control; W = 136, r = .68).

#### Produced sequence length reflects chunk-level forgetting

Even when imperfect, participants’ reproductions were not random. In particular, the distribution of the length of the produced sequence was strikingly structured (Fig B and Fig D in S1 Text; see Tables K and L in S1 Text for descriptive and centrality statistics). For Rep-2 (chunk size 2, repeated 6 times) and Rep-3 (chunk size 3, repeated 4 times), error lengths clustered around 10 and 9 respectively: precisely what would result from reproducing one fewer repetition than required (5 instead of 6, or 3 instead of 4). For Rep-4 (chunk size 4, repeated only 3 times), this effect was weaker. This pattern is consistent with Weber’s law applied to the mental representation of repetition count: the numbers 5 and 6 are perceptually close on a logarithmic scale, making a one-unit undercount likely while confusing 3 repetitions with 2 is less likely. In contrast, control sequences showed broad, much flatter error distributions with no such peaks, consistent with the lack of repetition structure anchoring a compressed mental representation. These patterns suggest that participants encode structured sequences as a count of repeated components, and that forgetting operates at the level of that count rather than at the level of individual items.

### Discussion

**Experiment 1** reveals that although memory for spatial locations is nearly perfect (virtually no token error), sequence reproduction depends heavily on sequence regularities, offering clear evidence that order-based regularity serves as a primary driver of working memory encoding. Specifically, structured sequences (repetitive and nested) significantly outperformed control sequences in both accuracy and production speed. Furthermore, while the classical short-term memory literature highlights both primacy and recency effects [29,30], **Experiment 1** was characterized primarily by a robust primacy effect. This early advantage is best understood through the lens of retroactive interference [31]: in control sequences, incoming items appeared to disrupt previous memory traces, whereas order-based structure specifically shielded the encoding of initial items. In contrast to the stochastic length errors of control sequences, structured sequences show systematic, block-wise omissions. Omissions in Rep-2 (peaking at 10 items) and Rep-3 (9 items) reveal that participants group items into bound subcomponents rather than storing 12 isolated units. This finding aligns with the subitizing and groupitizing phenomena [32–34] where quantity estimation is driven by group count rather than item count.

While **Experiment 1** qualitatively demonstrates the impact of order-based structure on memory, it does not provide a formal metric to quantify the complexity of the underlying mental programs or predict the specific code the brain employs. To move beyond mere observation, **Experiment 2** aimed to identify the unique minimal description for each sequence, by implementing a parameter-fitting approach on the costs of specific primitives. Specifically, we sought to determine if participants’ performance correlates with the LoT-complexity and if the “breakpoints” in the sequence descriptions align with the timing latencies (ICIs) and the constituent-based error patterns (such as the omission of entire blocks) that characterized participant performance.

Furthermore, **Experiment 1** focused exclusively on the repetition primitive as a primary driver of memory compression. However, the human capacity for sequence learning likely leverages a much richer repertoire of abstract regularities. To test the breadth of this mental “alphabet,” we expanded the stimulus set in **Experiment 2** to include more sophisticated operations, such as temporal inversion (Mirroring) and interleaving groups. This allowed us to investigate whether the human LoT is a flexible, multi-primitive system capable of exploiting diverse temporal structures to maximize memory efficiency.

## Experiment 2

Building on the behavioral evidence for hierarchical encoding in **Experiment 1**, **Experiment 2** sought to formalize these mental programs through a quantitative, predictive model. While **Experiment 1** established that order-based regularity facilitates memory, it did not quantify the specific “cost” of each mental operation involved. To translate these qualitative observations into an algorithmic framework, we adapted the Language of Thought (LoT) model, previously validated in the domains of geometry and binary sounds, to the study of abstract order-based sequences. To do so, we replaced the Rotation(n) primitive defined for the Language of Geometry [14–16,18,19] with a Move&Print(n) primitive, which represents movement within a vocabulary of ordinal positions.

The primary objective of **Experiment 2A** was to empirically determine the weights (costs) of these new core primitives. We assumed that repetitions and their variations share a common cost structure, defined by the logarithmic function *W_Repeat_*_(*n*)_ = *β* ∗*log* (*n*) + *γ*, with n the number of repetitions. This logarithmic parameterization reflects the information-theoretic cost of encoding an integer n in a minimal program (requiring log₂(n) bits). Similarly, we hypothesized that the cost of the Move&Print(n) primitive is independent of temporal direction (where n > 0 denotes the future and n < 0 the past). This weight was parameterized as *W_Move_*__*and*_*Print*(*n*)_ = 1 + *α* ∗ |*n*|. Where n denotes the ordinal distance between consecutive items (1-2, n=1; 1-3, n=2…). While earlier frameworks utilized arbitrary fixed costs, this parametrization allows us to account for the cognitive load increases with higher repetition counts or larger “ordinal distances”. We performed an exhaustive grid search on the behavioral data of **Experiment 1** to determine the optimal parameters. This refined model was then validated against new data from **Experiment 2A**. We hypothesized that this calibrated order-based LoT model would outperform standard complexity metrics in predicting both participant accuracy and reproduction timing patterns. The latter provides a window into the internal representation of sequences, allowing for a direct comparison with the model’s algorithmic sequence descriptions.

In **Experiment 2B,** we aimed to test whether the human encoding of new sequences calls for an extension of the proposed language, with additional operations serving as fundamental primitives. To do so, we introduced a set of exploratory sequences, each testing a particular rule. By comparing performance on these structured sequences against “control” versions (where the regularity was disrupted by a single item swap), we sought to determine which of these rules the human brain spontaneously uses for memory compression. If a rule is a genuine LoT primitive, participants should show significantly higher accuracy and distinct temporal patterns compared to the unstructured controls.

### Methods for Experiment 2A and 2B

#### Participants

The recruitment process, browser requirements, and ethical protocols were identical to those described in **Experiment 1**. 90 participants were recruited voluntarily and remained unpaid. Exclusion criteria were as follows: (1) total duration was an outlier (twice the mean duration); (2) all responses contained token errors. Based on these criteria, we excluded 2 participants. Our final dataset included 88 unique participants. Average duration of the experiment was 15 mins 38 secs.

#### Design and Procedure

The experimental design followed the same “delayed reproduction with betting” paradigm used in **Experiment 1**, with two primary modifications to the stimuli and timing. First, the stimulus duration on screen was increased from 200ms to 300ms to ensure robust encoding of the more varied sequences. Second, the stimulus set was expanded: while **Experiment 1** utilized 9 distinct sequences, **Experiment 2** included these original 9 (**Experiment 2A**) plus 16 new sequences (**Experiment 2B**), for a total of 25 unique patterns. Each sequence was presented twice, resulting in a total of 50 trials per participant.

#### Model Comparison

Previous work investigating LoT in sequences [15,16,18,19] typically assumes the number of repetitions or the distance involved in the rotation to have no impact on the cost of the operation. To address this limitation, we compared this baseline LoT model, which uses exactly the same parameter values as previous work on geometrical sequences [16], with two parametrizations of the order-based LoT model in which primitive weights are fitted to the data (Model 1 and 2). The full specification of the language, its primitives and the definition of LoT-complexity are given in S1 Text.

**Baseline LoT model**: Fixed weights *W_Repeat_*_(*n*)_ = 1, *W_Move_*__*and*_*Print*(*n*)_ = *W_Rotation_*_(*n*)_ = 2

**Model 1**: *W_Repeat_*_(*n*)_ = 1 + *β* ∗ log_2_(*n*) , *W_Move_*__*and*_*Print*(*n*)_ = 1 + *α* ∗ |*n*|

**Model 2**: *W_Repeat_*_(*n*)_ = *β* ∗ log_2_(*n*) + *γ*, *W_Move_*__*and*_*Print*(*n*)_ = 1 + *α* ∗ |*n*|

#### Parameters Search

We fitted 2 models by exhaustive grid search over the data of **Experiment 1**, one with 2 parameters: *α* and *β* and one with 3 parameters: *α*, *β* and *γ*. Each fitted cost took the values 0, 0.1, …, 5, that is 2,601 grid points for the two-parameter model and 132,651 for the three-parameter model, and we retained the costs of the grid point with the lowest AIC, assessed using linear mixed-effects models (R 4.6.0, lme4 2.0.6; [35]). As the plausible range of costs we report the set of grid points within 2 AIC of the best. We modeled edit distance as a function of complexity following the equation: *editDistance* ∼ *Complexity* + (1|*Participant*) where participant ID was included as a random intercept to account for global inter-individual variability. Models were compared by AIC, counting four parameters (intercept, slope, participant variance and residual variance). To prevent overfitting and to evaluate generalizability, we assessed the model’s performance using the parameter set identified on **Experiment 1** on data from **Experiment 2**, which involves a distinct participant sample.

#### Comparing LoT-structural and motor predictors of inter-click intervals

To assess whether inter-click intervals (ICIs) reflect the hierarchical structure predicted by the order-based LoT model beyond mere motor effects, we constructed two binary predictors for each inter-item transition (positions 1–11) of each sequence. The motor predictor encoded whether two consecutive items shared the same spatial position (1 = same position, 0 = different position). When the same position is repeated consecutively, shorter ICIs are expected due to reduced motor preparation demands, independently of any cognitive structure. The structural predictor encoded whether each transition crossed a constituent boundary as defined by the order-based LoT model’s minimal description (1 = boundary transition, 0 = within-constituent transition). Effective mental compression of sequences through the LoT would predict longer ICIs at constituent boundaries.

This analysis was restricted to the six sequences for which the two predictors diverged, namely Rep-2, Rep-3, Rep-4, Rep-Nested, Rep-Local, and Rep-Global. For all other sequences, motor and structural predictors were collinear, making their contributions impossible to dissociate. Only trials with perfect reproduction were included, to ensure that ICIs reflected intended, structured motor sequences rather than error-recovery processes. Data from **Experiments 1 and 2** were pooled, yielding one ICI observation per inter-item transition, per trial, per participant. Two linear mixed-effects models were fitted in R (lme4 [35]), both including a random intercept for participant to account for inter-individual baseline differences in response speed:

Model 1 (motor only): *ICI* ∼ *motor*_*predictor* + (1 | *participant*_*ID*)

Model 2 (motor and structure): *ICI* ∼ *motor*_*predictor* + *structure*_*predictor* + (1 | *participant*_*ID*)

Models were compared using a likelihood ratio test (χ² test via anova function).

#### Alternative complexity models

The following models were considered to compare with LoT.

##### Chunk Complexity

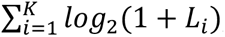 *log*_2_(1 + *L_i_*), with K the number of chunks and *L_i_* the size of the i^th^ chunk [9]. Chunk complexity depends heavily on how units are defined. We implemented two model variations: the Local model (based on Planton et al. [18]), which defines chunks as consecutive repetitions of single items (e.g., ‘11’) but is largely restricted to binary sequences; and the Global model, which defines a chunk as the minimal non-self-repeating unit that tiles an entire sequence (e.g., ‘123’ in ‘123123’).

##### Lempel-Ziv

Size of a dictionary built by a parsing algorithm [36]. The algorithm reads the string from left to right and adds to its dictionary every substring upon first encounter.

##### Shannon Entropy of Bigrams

*H* = − ∑*_b_ p_b_* ∗ *log*_2_(*p_b_*), with *p_b_* the relative frequency of bigram b among the 11 pairs of consecutive items of the sequence [37]. Following Planton et al. [18], entropy was computed over bigrams rather than single items, to capture first-order transitions; it is low when a few bigrams recur throughout the sequence.

##### Shannon Entropy of Transition Probabilities

*H* = − ∑*_i_* Σⱼ *A_i_*_j_ ∗ *log*_2_(*A_i_*_j_) with *A_i_*_j_ the empirical probability that item i is immediately followed by item j [37]. Unlike bigram entropy, this metric quantifies the unpredictability of the next item given the current one.

##### Shannon Entropy of L-HAL Associations

Same as above, but computed on the association matrix Â = (1 − *e*^−*β*^)*A(I* − *e*^−*β*^*A)*⁻¹ of the long-horizon associative learning (L-HAL) model [38], with I the identity matrix. In L-HAL, each item is also associated with more distant items, with a strength that decays exponentially with distance. As in [38], β was fixed to 0.06 rather than fitted to our data.

##### Subsymmetries

Mirror-based structural redundancies between substrings [39]. Although our repetition-based sequences lack such symmetries, the model was included for completeness and comparison with alternative models.

##### Compression Complexity

This metric approximates the Kolmogorov-Chaitin complexity of a sequence, i.e., the length of its shortest generating program [40,41], by its size after compression [42–44] with the zlib DEFLATE algorithm, with complexity values derived from the resulting byte length; lower values indicate higher compressibility driven by multi-level local repetitions and structure. Like LoT-complexity, this measure is thus a description length relative to a particular language, here that of the DEFLATE algorithm rather than the order-based LoT model.

##### Change Complexity

This metric measures the amount of variation across all substrings in a sequence and has been shown to align well with human performance across various cognitive tasks [45].

##### Subjective Complexity

Mean subjective complexity rating of each sequence (on a scale from 1 to 7, see **Experiment 3**).

#### Sequences tested in Experiment 2B

**Experiment 2** included the original 9 sequences of **Experiment 1** plus 16 new sequences. These new sequences were designed to determine if other operations should be added as primitives to the order-based LoT model to correctly account for humans’ sensitivity to sequence regularities (see Table 1 for stimuli sequences expressions). Each of these sequences was constructed by hand to instantiate one candidate operation, together with a control in which this regularity is specifically disrupted by a minimal change (see Table 1).

**Table 1.**
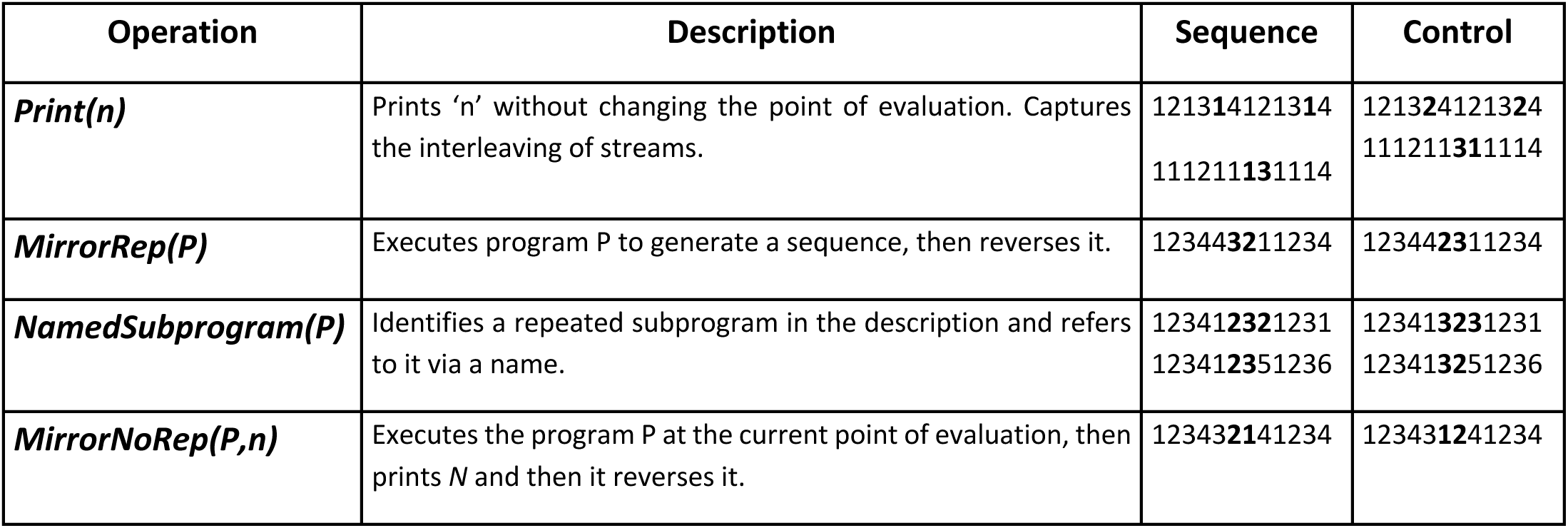
Candidate LoT primitives: structured sequences and minimal controls for empirical validation. This table presents structured sequences generated using a new set of candidate primitives (left), alongside minimally modified control sequences (right), used in **Experiment 2**. These sequences were designed to evaluate whether new operations should be incorporated into the order-based LoT model. Control sequences were constructed by introducing minimal changes at constituents’ boundaries, disrupting the original structure while preserving surface similarity (edit distance between original and control sequences ranges from 1 to 2 across all pairs). Differences are marked in bold and underlined.

**Table 2.** Summary of candidate rules tested as potential LoT primitives. A rule is retained if the structured sequence shows a significantly lower edit distance than its control. Additional criteria include evidence of retroactive memory facilitation and interclick time patterns consistent with the proposed structure. Asterisks indicate significance levels (* p < .05, ** p < .01, *** p < .001). The final column shows the LoT-complexity under the order-based LoT model (i.e., without the candidate primitives) for the structured/control sequences.

| Operation | Retained | Edit distance difference | Retroactive facilitation | Interclick times structured by rule | LoT-Complexity structure / control |
| --- | --- | --- | --- | --- | --- |
| Mirror-Rep | Yes | Yes*** | Yes*** | Yes*** | 16 / 19 |
| Mirror-NoRep | No | No | No | N/A | 20 / 24 |
| NamedSubprogram-1 | Yes | Yes*** | Yes* | N/A | 21 / 22 |
| NamedSubprogram-2 |  | Yes*** | Yes** | Yes* | 23 / 27 |
| Print | Yes | Yes*** | No | Yes*** | 21 / 21 |
| Print-4 |  | Yes*** | Yes*** | N/A | 18 / 13 |

##### New primitives

*Print* calls a selected element without moving the point of evaluation. For instance, in the tested sequence 121314121314, the point of evaluation advances through 1→2→1→3→1→4 (via *Move&Print*), while ‘1’ is repeatedly “printed” between each step without advancing. This primitive was introduced to capture the interleaving of two streams: a stable stream (e.g., repeated 1’s) and a varying stream (e.g., *2, 3, 4*). The sequences referred to as “Print” and “Print-4” were designed to test this primitive.

*Mirror* temporally reverses a group of elements. To determine if repeating the last element is necessary to the *Mirror* primitive, we separated *Mirror* into two proposed versions. The *Mirror-NoRep* version is a *Mirror* without repetition of the last element of the group. *MirrorRep* repeats the last element of the group on which it applies (see Table 1 for the sequence implementation). NamedSubprogram stores a subsequence under a reusable label (e.g., ‘123’) and re-invokes it at multiple points in the sequence, allowing the same motif to recur even when separated by intervening items (e.g., 123-4-123-5-123-6). This structure reflects the idea that reusing a previously defined subprogram should reduce complexity. In the control conditions, this regularity is disrupted by altering one of the repeated subsequences, thereby interfering with the identification and reuse of the common pattern.

#### ICI slow down

Inter-click intervals were computed on cumulatively correct responses: an interval at position *i* was included if all items up to and including position *i* had been correctly reproduced, regardless of accuracy at later positions. This preserves intervals from partially correct trials, in which the participant’s reproduction had not yet diverged from the target.

### Experiment 2A Results: Model Calibration and Validation

#### LoT outperforms other complexity models

Building on previous frameworks [16,46], we refined the weights for the Move&Print(n) and Repeat(n, P) on **Experiment 1** data and validated them against **Experiment 2** data, collected on a separate population sample. The fitted costs were α = 1.0 and β = 1.0, so that repeating a pattern k times costs 1 + log₂(k), exactly the number of bits needed to encode k. Two regions of the cost space fitted **Experiment 1** almost equally well. The first lies around α = β = 1. The second runs along the line β = α + 1, with α from 0.8 to 1.0 and β from 1.4 to 1.9, where doubling the number of repetitions of a block costs as much as one additional move of one ordinal step. Its best point (α = 0.8, β = 1.8) was only 1.15 AIC above the best grid point, so both regions fell within the plausible range of costs. The two generalized to **Experiment 2**almost equally well (AIC = 6149 vs. 6151; r = .958 vs. .957, difference in r = 0.001, 95% CI [- 0.005, 0.007]). We retained α = β = 1, the best grid point on **Experiment 1**. Once the additional fitted cost is counted, the two- and three-parameter models were equivalent on **Experiment 1** (AIC = 3418.93 and 3418.51), but the two-parameter model generalized better to **Experiment 2** (AIC = 6149 vs. 6162, ΔAIC = 13). We therefore retained the two-parameter model. Nothing was refitted on **Experiment 2**, and the high Pearson correlation obtained there (r = .958, 95% CI [.937, .970], p < .001) between edit distance and LoT-complexity (**Fig 2B**) confirms the model’s predictive strength for behavioral data. To assess how LoT-complexity compares to previously established complexity metrics, we evaluated the explanatory power of each measure using mixed-effects models. In these models, alternative complexities were treated as a fixed effect, while participants were included as a random effect to account for inter-individual variability in baseline performance.

Across all models tested (see **Fig 2C**), LoT-complexity consistently provided the best fit (lowest AIC) to participant performance, replicating prior findings in the visual domain and highlighting its robustness as a framework for capturing how humans structure sequences in memory [15,18].

#### Sequence descriptions obtained from the order-based LoT model

The minimal programs of **Fig 3** were derived from the model. Their length immediately explains why LoT is a good predictor of behavior: while the structured sequences used hierarchical nesting and recursion to achieve short description lengths, the control sequences remained long and fragmented.

**Figure 3.**
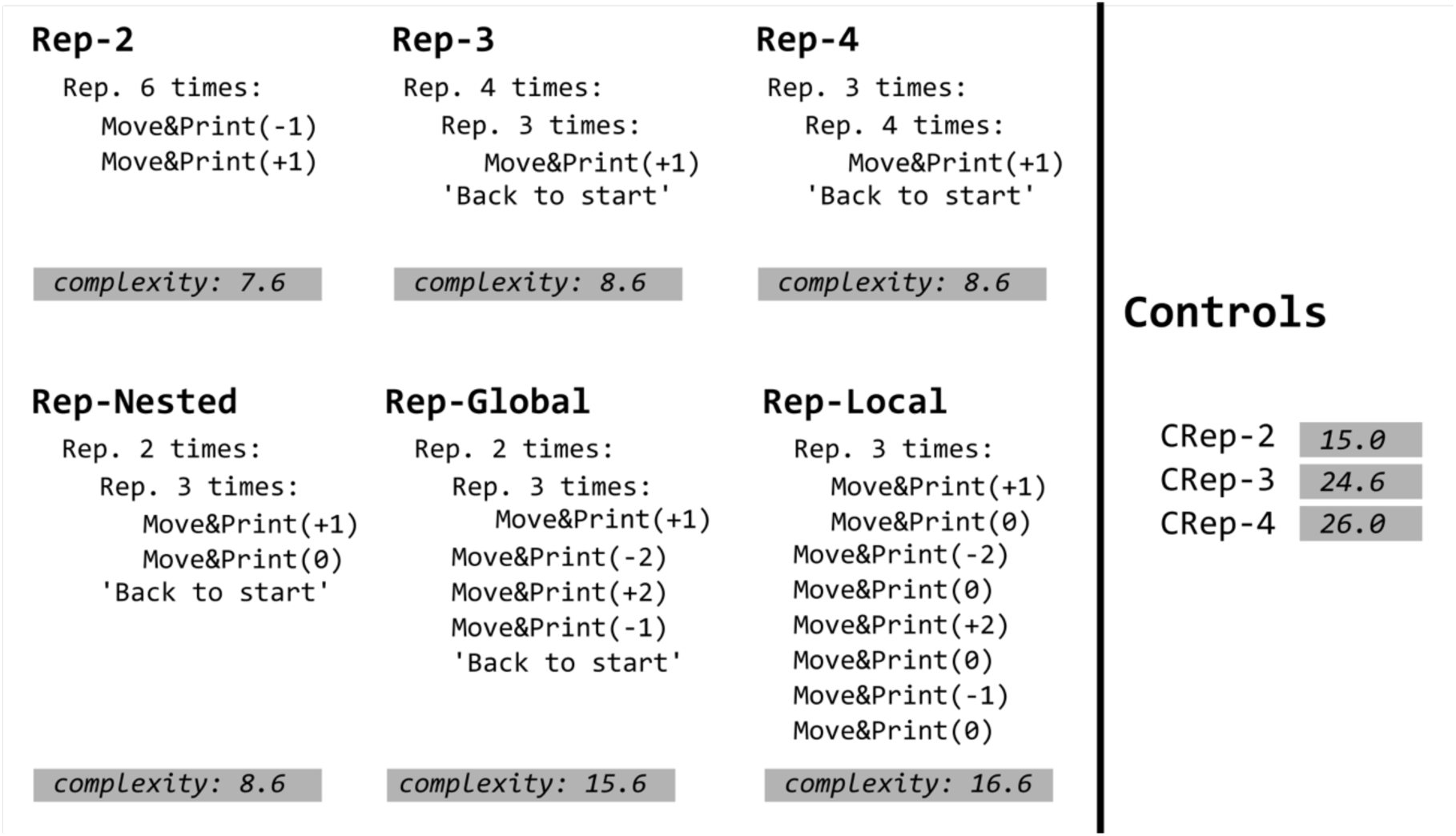
LoT programs and complexity for repetition-based sequences used in Experiment 1. For each sequence, the minimal program inferred by the order-based LoT model is shown, together with its LoT-complexity (minimum description length with weighted primitives). Programs are composed of the primitives defined in **Fig 1A** and in S1 Text. “Back to start” is a special case of “Repeat n times with application of ‘instruction’”, with instruction being “Move&Print(0)”. “Back to start” returns the point of evaluation to its initial position. Structured sequences (Rep-2, -3, -4, and -Nested) are described by short programs exploiting nested repetition. Rep-Global and Rep-Local require substantially longer descriptions and serve as controls isolating hierarchical level for Rep-Nested. Complexity values for the three matched controls (CRep-2, -3, -4) are shown at right; these sequences use the same elements as their structured counterparts but in a rearranged order, and resulted in unstructured, long programs.

#### Sequence structure impacts on participants’ reproduction timing

We examined whether perceived structure, measured by inter-click intervals (ICIs; measured on perfect sequence reproduction trials) and cumulative accuracies aligned with LoT-description constituent boundaries (see **Fig 4**). Increases in ICIs suggest motor preparation for new constituents, while drops in cumulative accuracy typically mark constituents’ boundaries [47]. When two successive items appear at the same spatial location, shorter ICIs are expected compared to when items are spatially separated. To assess whether ICIs reflect the hierarchical structure of sequences beyond mere motor effects, we compared two linear mixed-effects models: one accounting for motor facilitation when the same item is repeated consecutively, and another including, as an additional predictor, the hierarchical structure derived from the LoT model (see *Methods*). Both models included a random intercept for participant. The hierarchical structure predictor significantly improved model fit compared to a model with only the motor predictor (χ²(1) = 107.64, *p* < .001; ΔAIC = 106), indicating that constituents implied by the hierarchical structure of sequence LoT description contribute significantly to ICIs variability.

**Figure 4.**
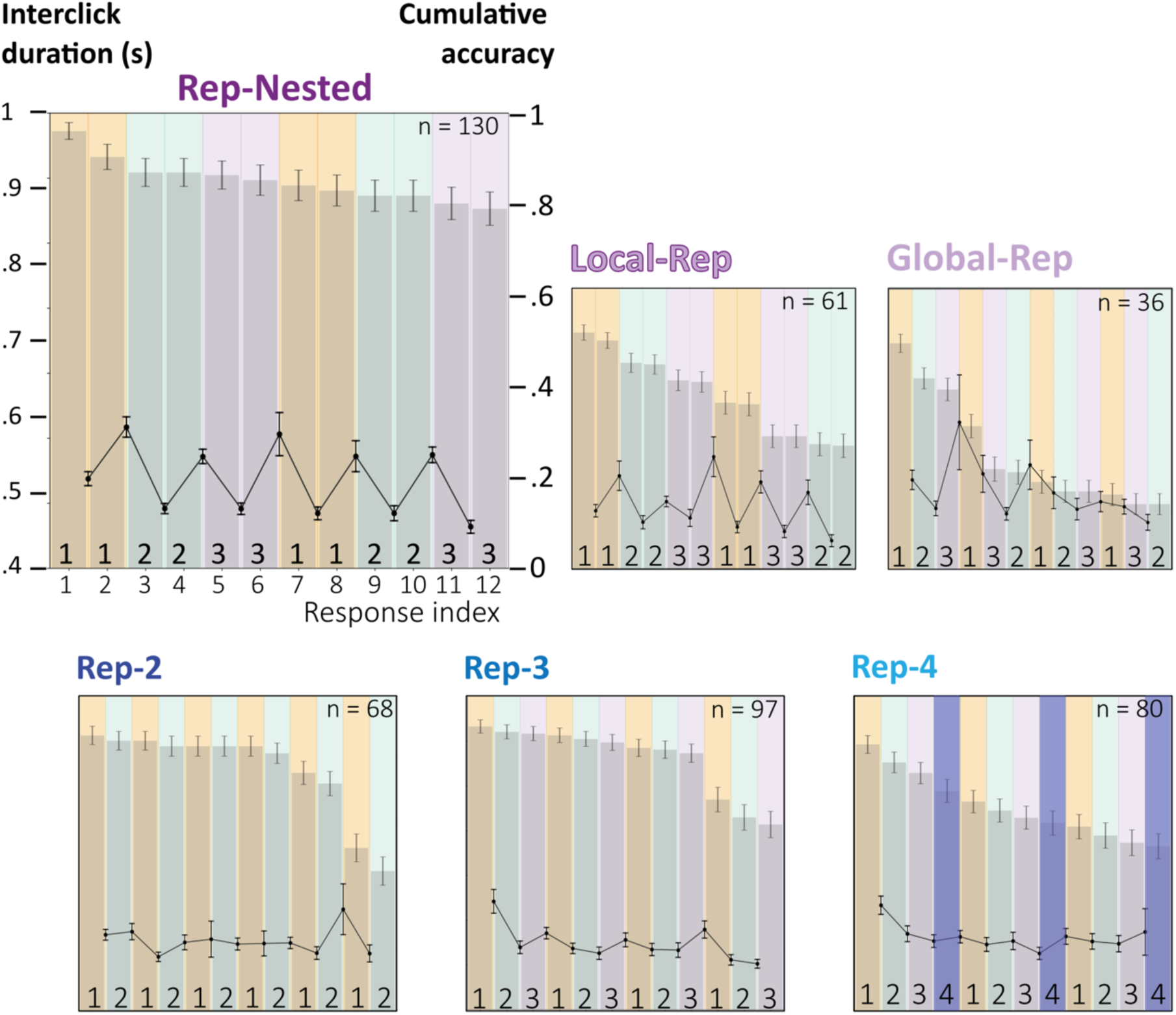
Behavioral markers of chunking align with temporal rules. The figure overlays (1) cumulative accuracy (background histograms) and (2) inter-click intervals computed on accurately reproduced sequences. Colors indicate sequence order-based structure (1: orange, 2: green, 3: gray, 4: purple). Error bars represent the standard error of the mean. The number of correct responses for each sequence (**Experiment 2** only) is displayed in the top-right corner of each panel. Drops in cumulative accuracy and peaks in inter-click durations mark chunk boundaries [47].

We further analyzed the behavioral markers of sequence structure for each sequence type separately. Across all sequence types, inter-click intervals (ICIs) at constituent boundaries were significantly higher than within-constituent transitions, indicating sensitivity to LoT structure (see **Fig 4**). In Rep-2 and Rep-3, boundary ICIs were elevated (Rep-2: 482 vs. 443 ms, p = .005, r = .41; Rep-3: 465 vs. 433 ms, p < .001, d = .44) and accuracy dropped sharply at the boundary of the final constituent (positions 10–11 and 9–10, respectively), suggesting constituent-level forgetting. Rep-4 showed a more gradual decline and no reliable boundary effect (472 vs. 454 ms, p = .455). In Rep-Nested, local repetition boundaries drove clear ICIs (485 vs. 395 ms, p < .001, r = .75), with no additional cost at the global boundary (p = .40). In Global-Rep, constituent boundaries were marked (590 vs. 472ms, p = .005, r = .52), but subcomponent and global boundary intervals did not differ (p = .73), and accuracy dropped mainly at positions 3–5. In Local-Rep, boundary ICIs were elevated (498 vs. 406 ms, p < .001, r = .74), with a distinct peak at the sequence midpoint (560 vs. 402 ms, p < .001, r = .81), consistent with accuracy drops at positions 6–7 and 8–9. All of these effects replicated in the **Experiment 1** dataset, with closely matching effect sizes.

### Experiment 2B Results: Sequences challenging the order-based LoT model

#### Which regularities facilitate sequence reproduction?

To assess whether a candidate primitive improves the order-based LoT model (Fig 5A), we compared the initial model to an extended model that also included each candidate primitive predictor (coded 0 for primitive sequences and 1 for controls). The comparison aimed to assess whether the candidate primitive improved compression in a test sequence relative to its control. This comparison yielded significant model improvement for *Mirror-Rep* (ΔAIC = -66 and p < .001) and *Print* (ΔAIC = -29 and p < .001). No significant difference was found for *Mirror-NoRep* and *NamedSubprogram*. To corroborate the potential inclusion of new operations into the LoT, we examined whether behavioral markers, specifically inter-click intervals (ICIs) and error patterns, aligned with the hypothesized rules.

**Figure 5.**
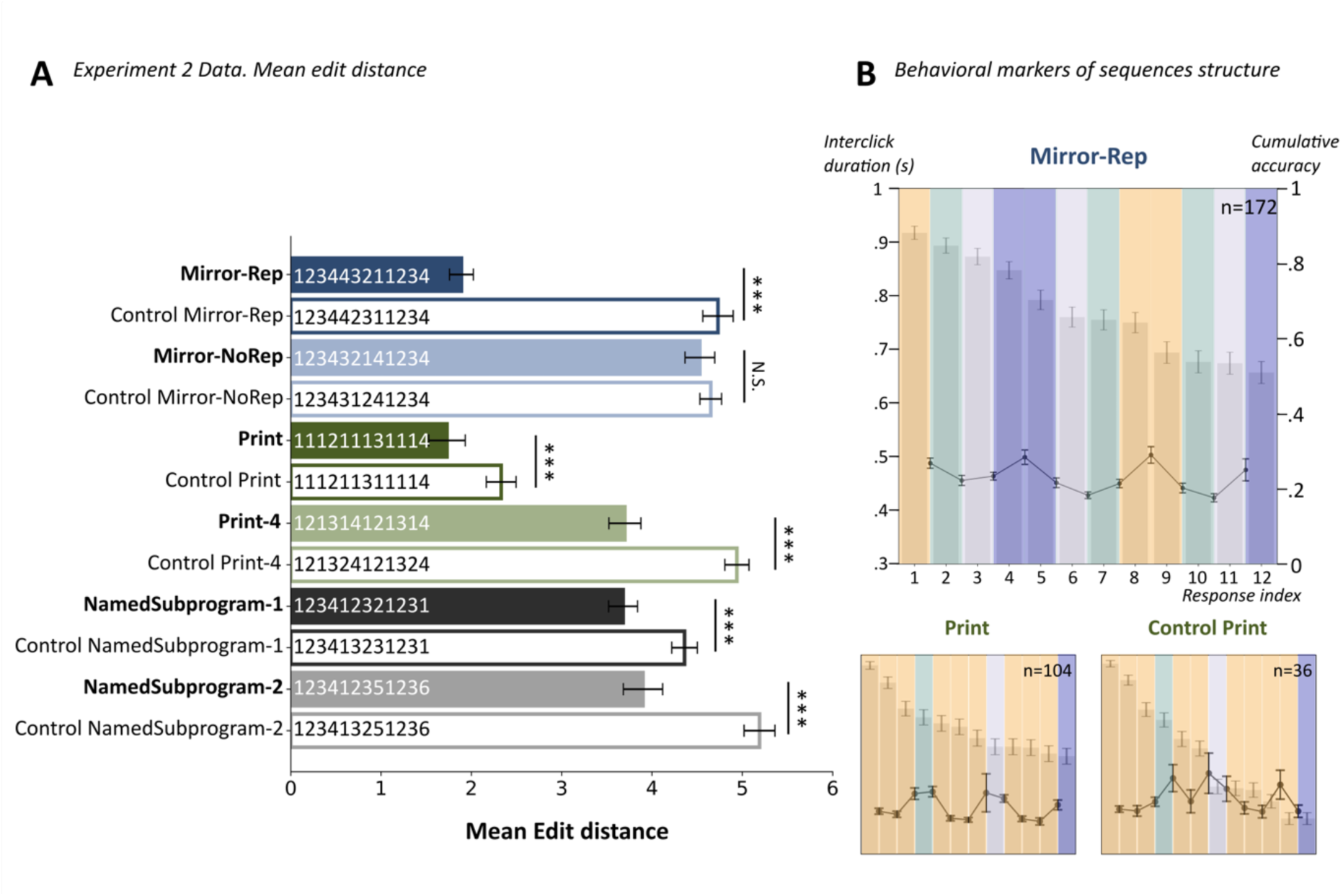
Performance and behavioral markers of chunking reveal additional temporal primitives in sequence recall. (A) Performance, measured as mean edit distance, shows that exploratory primitives may structure sequence memory. Asterisks denote significance in pairwise comparisons of per-participant mean edit distance, assessed with a Wilcoxon signed-rank test (n.s. p > .05, * p < .05, ** p < .01, *** p < .001). (B) Behavioral markers of chunking show that chunk size and boundaries are defined by temporal rules. Panels overlay (1) cumulative accuracy (background histograms) and (2) inter-click intervals computed on accurately reproduced sequences. Colors indicate sequence order-based structure (1: orange, 2: green, 3: gray, 4: purple). Error bars represent the standard error of the mean. The number of correct responses for each sequence (**Experiment 2**) is displayed in the top-right corner of each panel. Drops in cumulative accuracy and peaks in inter-click durations mark chunk boundaries [47].

##### MirrorRep

ICIs were significantly longer at mirror boundaries (‘1234—4321’; 503ms vs. 441ms, *p* = .006, r = .41, see **Fig 5B**), and cumulative accuracy dropped notably at those same points (between items 4–5 and 8–9) despite the presence of local repetitions at these points. This suggests that the mental chunking followed the global mirror symmetry rather than the local repetitions (‘44’) predicted by the order-based LoT model (Model 1, see Results Section from **Experiment 2A**; see Table A in S1 Text; see Fig C in S1 Text for cumulative accuracy and mean ICIs).

##### NamedSubprogram

In the NamedSubprogram-2 sequence, elevated ICIs at the 1234–1235–1236 boundaries (499 ms vs. 455ms, p = .02, r = .41) and a sharp accuracy drop at the 5^th^ item suggest that participants utilized a subprogram template. This segmentation differed significantly from the one expected by the order-based LoT model.

##### Print

For the Print-4 sequence, but not the Print sequence, early-item accuracy showed retroactive interference (p < .001; see Table N in S1 Text). For Print, ICIs were significantly higher at chunk boundaries (507ms vs. 456ms, *p* < .001, r = .61), again diverging from the order-based LoT model description.

ICIs’ alignment with the rule could not be assessed when not enough correct trials were available for analysis. It was the case for Print-4 (error rate: 85.83%, SEM = 2.65%), NamedSubprogram-1 (94.96%, SEM = 1.5%) and Mirror-NoRep (97.01%, SEM = 1.39%).

Together, Mirror-Rep, NamedSubprogram, and Print emerged as strong candidates for inclusion in a revised order-based LoT model. Conversely, Mirror-NoRep could be confidently excluded as it lacked clear behavioral advantages.

### Discussion of Experiment 2

The primary objective of **Experiment 2A** was to characterize the order-based LoT model by calibrating primitive costs and validating its predictive power against independent data. The optimized model achieved high correlation with behavioral performance (r = .96), significantly outperforming traditional metrics like Shannon entropy or Lempel-Ziv. Critically, the model’s utility extended beyond a general metric of performance (edit distance); it accurately predicted the fine-grained temporal dynamics of sequence reproduction. The observed inter-click intervals (ICIs) provided direct behavioral evidence that participants encode sequences as nested programs rather than linear chains. Increases of ICIs at constituent boundaries suggested that the brain incurs a quantifiable cognitive cost when transitioning between mental sub-programs, a signature that is captured by the LoT’s structural description.

**Experiment 2B** demonstrated that the human LoT is more complex than the order-based LoT model that we used so far and required a broader set of primitives to fully capture the entire range of human sequence compression abilities. Our findings indicated that Mirror-Rep (temporal symmetry) and Print (interleaving) are strong candidates for core primitives. The behavioral signatures (that is, enhanced performance and boundary-specific ICIs) suggested that these operations are integral to the adult human representational system. Conversely, more abstract or computationally demanding operations like NamedSubprogram and Mirror-NoRep yielded inconclusive results. This suggests that not all formal regularities are exploited with equal facility. Future research will be needed to determine whether these primitives generalize across different sensory modalities (e.g., geometric, auditory) and across development.

If the order-based LoT model is very good at predicting objective behavior, are participants explicitly aware of the underlying “code“? Our results suggest that this LoT operates as a pre-existing representational system that does not require the “discovery” or learning of new primitives during the task. Instead, participants appear to dynamically assemble primitive operations on the fly to achieve real-time compression of the sequences. As a consequence, the underlying computational mechanics may bypass explicit monitoring. **Experiment 3** investigated this potential metacognitive opacity by contrasting objective LoT-complexity with subjective ratings, determining whether the metacognition maintains an accurate map of its own computational constraints or if the LoT machinery remains introspectively inaccessible.

## Experiment 3. Subjective complexity

### Methods

Eighty participants were recruited via Prolific.com. Participants were paid at a rate of £9 per hour, in line with Prolific’s payment guidelines (approximately £2.50 per participant). Anonymization, browser restrictions and the consent procedure were identical to **Experiment 1**, with an information page adapted to this experiment; it was approved by the same committee under amendment CER-Paris-Saclay-2019-063/A3. The 25 sequences from **Experiment 2** were each presented 3 times (75 trials total). The presentation phase was identical to **Experiment 2**, followed by a subjective complexity rating: participants answered “How difficult was that sequence to memorize?” on a 7-point Likert scale (1 = Very Simple, 7 = Very Complex). A short training session with three anchor sequences calibrated participants’ use of the scale. No feedback was provided. Probe sequences (one easy: ‘111111111111’; five difficult) were used to exclude inattentive participants (criteria: rating > 2 on easy probes or < 3 on difficult probes), leading to the exclusion of 3 participants (final N = 77; median duration = 16.5 minutes).

#### Model fitting and comparison

Models were fitted following the same procedure as in **Experiment 2** (linear mixed-effects models, lme4, AIC-based comparison, participant as random intercept). To benchmark subjective complexity against LoT complexity, we compared their respective AIC values on the same sequences. Pearson and Spearman correlations were computed between subjective complexity ratings, LoT complexity scores, and behavioral performance.

### Results

Probe responses confirmed attentiveness: the easy probe was rated near floor (mean = 1.15, SEM = 0.04), while the five difficult probes received consistently high ratings (means 5.05–5.64). Ratings were distributed across the full scale (mean = 3.80, SD = 1.85), with no floor or ceiling concentration. Mean subjective complexity ratings correlated strongly with reproduction accuracy across the 25 sequences (r (23) = .89, p < .001; Spearman ρ = .83, p < .001). Nevertheless, LoT complexity provided a substantially better fit to the behavioral data than subjective complexity in mixed-effects model comparison (ΔAIC = 120.77; LoT: AIC = 6148.68 vs. subjective: AIC = 6269.45).

### Discussion

**Experiment 3** showed that subjective complexity judgments track LoT complexity (r = .89), indicating that participants have reliable conscious access to the order-based structure of sequences, which is consistent with findings from the geometrical domain [48]. Nevertheless, LoT complexity provided a substantially better account of reproduction accuracy (ΔAIC = 120.77), suggesting that the LoT captures structures that influence memory encoding beyond what participants can explicitly report. Subjective complexity thus appears to be a valid but coarse proxy for LoT complexity, informative about perceived difficulty, but insufficient as a standalone predictor of memory performance.

## General Discussion

Working memory for structured sequences is determined by the length and structure of the mental program that describes them. Previous work successfully established this principle to visuo-spatial sequences governed by geometric rules and binary auditory sequences with regularities. Here, we show that the same principle applies to regularities in a more abstract domain, namely ordinal memory, in which items are coded by their order of first appearance rather than by their physical identity. To do so, we implemented an order-based Language of Thought (LoT) model and tested it in a series of memory tasks. Minimum description length (LoT-complexity) outperformed other sequence complexity metrics in predicting memory performance, and also outperformed participants’ subjective complexity judgments. Beyond accuracy, programs inferred by the model aligned with behavioral markers of motor execution’s structure: inter-click intervals slowed down and errors increased more sharply at the constituent boundaries the model specified. Together, these results indicate that when a sequence affords a discoverable order-based structure, the human brain encodes it in working memory as a mental program. It then determines both retrieval performance and the timings of motor productions, yet the program remains only partially accessible to introspection. The compositional principles previously identified in the visuo-spatial and auditory domains therefore govern the more abstract order-based structure [14–16,18,19].

### A Language of Thought for order-based structure

Our experiments establish that an order-based LoT model, built on primitives like repetition (*Repeat*) and relative movement in ordinal space (*Move&Print*), accurately predicts how humans encode structured sequences. The model’s complexity metric showed a remarkably strong correlation with memory performance (r = .96) and significantly outperformed a wide range of alternative complexity models, including those based on chunking, Shannon entropy, and compression complexity. Crucially, the mental abstract representations proposed by our model have concrete behavioral signatures: participants’ response timings revealed a hierarchical constituent structure that aligned with the boundaries predicted by the LoT’s most compressed descriptions. This demonstrates that humans spontaneously parse temporal information into nested, rule-based components. One feature of the fit deserves emphasis. The weight of the repetition count was free to take any value. The best fit put it at exactly one. The instruction itself costs one unit, so repeating a pattern k times costs 1 + log₂(k). Its argument is then charged precisely the number of bits needed to encode k. The data do not pin this value down completely: a second region of the cost space, with a repetition weight near two, fits **Experiment 1** almost as well. But the best fit falls exactly where the cost structure of the mental language tracks the information content of what it encodes, rather than an arbitrary psychological scale.

Furthermore, we showed that this mental language is not limited to simple repetition. In our second experiment, participants successfully leveraged more sophisticated rules, such as temporal inversion (*Mirror*) and the interleaving of streams (*Print*), to improve their recall. The discovery of these additional primitives suggests that the human mind possesses a rich, extensible vocabulary for describing temporal patterns.

### Compression without introspective access

It is notable that participants’ subjective complexity judgments were outperformed by the LoT-complexity metric. Several factors may explain this discrepancy. The explicit judgment task (rating complexity on a scale) is fundamentally different from the implicit reproduction task, which directly engages compression mechanisms in working memory. Furthermore, it is likely that participants have only limited conscious access to the compression algorithms their brain executes. Thus, their intuition of complexity may not faithfully reflect the actual computational cost of encoding, which the LoT-complexity appears to capture more effectively.

### Sequence compression as a signature of human cognition

A growing body of evidence suggests that the human capacity for information compression thanks to minimal mental programs may be a defining signature of human cognition, realized through domain-specific languages of thought [17]. This hypothesis has gained traction across independent cognitive domains. In geometry, Sablé-Meyer and colleagues demonstrated that the perceived simplicity of a shape is predicted by its minimal description length in a generative language of geometric primitives [48,49]. In the domain of number, the frequency of number words in natural language reflects their compositional complexity within an algebraic LoT built on addition and multiplication [50]. Each of these findings establishes the same principle within a distinct representational system. Whether this capacity is specifically human is less obvious than it may seem. Humans do outperform other primates on structured sequences [51,52], yet much of the underlying machinery appears to be shared: monkeys are limited in serial recall and visual working memory in much the same way we are [53,54], they track sequential regularities through largely similar mechanisms [38,55–59], and neurons in the macaque frontal cortex code abstract roles and ordinal positions within a learned grammar [60]. What they master after extensive training nevertheless remains far simpler than the multi-level programs our participants assembled from a single presentation [61–63].

### Order as an abstract feature domain

Each mental language proposed so far operates on a specific kind of content. The geometric primitives move and trace in space [48,49], the arithmetic ones add and multiply quantities [50]. Order-based structure offers nothing of the sort to work on. The Language of Thought model encodes order-based structure as mental programs describing transitions between ordinal memory slots. Crucially, spatial positions were randomized across trials, and control sequences used the same items as structured ones, so that the compression advantage cannot be explained by geometry. The fact that the Language of Thought model predicts performance with high fidelity in this purely order-based setting (r = .96) suggests that program induction does not need to be tied to any specific sensory or spatial code, but can operate over abstract relational structure as such. The primitives we identify appear to function as ordinal operations for manipulating temporal regularities, just as Turn and Trace serve geometrical representation [16], or +/× serve number [64]. It remains untested whether some primitives from distinct domains map onto each other, recycling similar neural code, or whether each domain possesses a certain set of unique and specific primitives for LoT compression.

These behavioral principles may have a neural counterpart. In rodents, El-Gaby et al. [65] showed that neurons of the medial frontal cortex form structured memory buffers. These buffers tile progress toward a goal, irrespective of sensorimotor specifics. Individual cells fire at fixed lags relative to behavioral anchors. The result is a programmable scaffold that encodes a sequence through network dynamics alone, and that transfers to new goals without task-specific learning. In humans, Shpektor et al. [22] report a hierarchical ordinal code in the entorhinal cortex that locates an item within an auditory sequence using several factorized coordinates, organized in a nested hierarchy along the anterior-posterior axis. Both findings describe an ordinal code that is decoupled from elapsed time and from physical distance. Both organize it hierarchically. This is what our participants’ behavior requires. The primitives we identify, and their nesting, may therefore rest on a pre-wired architecture for abstract sequence memory, rather than on representations built anew for each task [65–68].

### Limits of the present account

#### What program length does not capture

Two sequences whose shortest programs have the same length are predicted to be equally difficult. This is not what we observe. Three sequences of **Experiment 1** receive exactly the same complexity (**Fig 3**), yet they were not reproduced equally well (**Fig 2B**). A central open question is how to score reproduction errors in a way that accounts for structure. Our edit distance measure, while representing an improvement over raw error rate or length of correct subsequence, remains insensitive to the sequence’s structure; it penalizes errors (e.g., a substitution or omission) uniformly, regardless of whether the error breaks a regularity. Our own results on cumulative performance show that errors are more likely to occur at the boundaries of structural chunks. As noted by Mathy & Varré [69], traditional global similarity metrics may obscure how structural regularities influence recall performance, motivating the search for more nuanced scoring methods.

#### Combining sources of structure

Our experiment was designed to minimize the role of geometry: spatial positions were assigned randomly, and control sequences used the same items as structured ones. This allowed us to isolate the encoding of order-based structure. However, this raises a crucial question for the future: how can the distinct LoT models for order-based and geometrical regularities be integrated into a single model of working memory for sequences with spatiotemporal regularities? Future work should investigate sequences where both types of structure are present and equally relevant, in order to model how the brain combines these different sources of information into a unified representation.

#### From a deterministic to a probabilistic language

While our order-based LoT model successfully captures a significant portion of the variance in human sequence memory, a natural next step is to add a noise model to the LoT. An example of this necessity is with the Approximate Number Sense, where the mental representation of magnitude becomes increasingly imprecise as the value grows: a principle governed by Weber’s law [70,71]. With a probabilistic framework, our model could predict errors such as a participant miscounting six repetitions as five, something yet unaccounted for.

An interesting case our framework does not cover yet is the hybrid sequence: “a simple rule, plus some exceptions”. The present fully deterministic order-based LoT model would generate an overly complex description for the random part. Future work should explore how our framework could be extended to allow for such hybrid representations, perhaps by integrating a mechanism that partitions a sequence into deterministic and stochastic components [72]. This would provide a more psychologically plausible account of how we handle partially predictable environments. Our approach, based on finding a single compressed description, contrasts with influential probabilistic frameworks that model learning as Bayesian inference over a space of generative rules or programs [73–75]. These models yield a probability distribution over possible structures rather than a single complexity score. Though methodologically different, both approaches frame learning as a search for simple, explanatory descriptions, a core principle in language and thought [11]. A powerful synthesis would use our compact LoT primitives within such a probabilistic framework, combining the strengths of interpretable rules and flexible Bayesian inference.

This line of work outlines a research programme with three complementary aims: identifying, for each type of structure, the primitives and composition rules that best account for human behavior; determining which of them are shared across domains, as expected if they reflect a general capacity for compositional compression rather than a collection of domain-specific tools; and comparing these mental languages across species, to test whether the capacity to nest and recombine primitives is a signature of human cognition [17,48,51,55,60]. Order-based structure offers a privileged test case for the second aim: because it depends only on the order in which items appear and repeat, and not on their nature, it can be carried by any modality. Consequently, this study offers a candidate representation that enables direct empirical testing of amodal structures.

## Conclusion

When a sequence afforded an order-based regularity, participants encoded it as a program, assembled on the fly from a small vocabulary of primitives: repeat, move and print, mirror, interleave. The length of that program predicted how well the sequence was recalled, and its structure showed in the timing of the taps. The compositional principle already established for geometric shapes, auditory patterns and number words therefore extends to structure carried by order alone. This is an abstract code that belongs to no sensory modality in particular.

## Supporting information

merged supplemental files

## Acknowledgments

Elyes Tabbane is a doctoral student in the FIRE PhD program, funded by the Bettencourt Schueller Foundation and the EURIP Graduate Program (ANR-17-EURE-0012). This research was conducted within the framework of Université Paris Cité and Neurospin (CEA Saclay).

This work was supported by INSERM (Institut National de la Santé et de la Recherche Médicale), CEA (Commissariat à l’Energie Atomique et aux Energies Alternatives), Collège de France and the Bettencourt Schueller Foundation. This work was also supported by a FYSSEN research Grant to Fosca Al Roumi, and a European Research Council ERC advanced grant ‘MathBrain’ to Stanislas Dehaene.

We gratefully acknowledge extensive discussions with Christophe Pallier, Fabien Mathy and Mathias Sablé-Meyer. We thank all the volunteers for their participation. We are grateful to Isabelle Denghien-Courcol for her help on deploying the online experiments.

## Availability statement

All behavioral data underlying the reported findings are deposited on Zenodo (https://doi.org/10.5281/zenodo.20070505). Analysis and model fitting code is available at https://github.com/ElyesT0/sequence_memory_ordinal_structure_analysis. The LoT model implementation is publicly available at https://github.com/sfigueir/lot-sequence-complexity (commit 4be6377).

## Supporting information

S1 Text. Supplementary methods, figures and tables. Formal specification of the order-based Language of Thought (machine state, primitives, concatenation, repetition constructs, program size and LoT-complexity, and the primitive weights used in **Experiment 2**), together with Figs A–D and Tables A–N.

