## Supplementary material for "Humans compress order-based structure into short mental programs": merged supplemental files

### The Language of Thought Programming Language

#### Machine state

The order-based Language of Thought (LoT) used in the paper operates over a cyclic set of $b$ ordinal memory slots,

$$\mathcal{A}_b = \{0, \dots, b-1\}.$$

At any moment, the state of the machine consists of:

(i) a *point of evaluation* $x \in \mathcal{A}_b$, indicating the current ordinal memory slot; and

(ii) an output sequence $s$.

A program starts from a specified point of evaluation and an empty output sequence. Instructions may change the point of evaluation, append symbols to the output, or both. All movements of the point of evaluation are computed modulo $b$.

The semantics of the instructions used in the paper are described below. For clarity, each description is accompanied by short Python-like pseudocode.

#### Move&Print$(n)$

The instruction

$$\text{Move\&Print}(n)$$

moves the point of evaluation by $n$ ordinal slots and outputs the item stored in the slot it reaches.

```
def move_and_print(n, position, base):
    position = (position + n) % base
    return str(position), position
```

For example, with $b = 6$,

```
Move&Print(+1)
Move&Print(+1)
Move&Print(0)
Move&Print(-2)
```

executed from $x = 0$ produces

$$1220.$$

The same program produces $2331$ from $x = 1$ and $5004$ from $x = 4$.

In particular, $\text{Move\&Print}(0)$ leaves the point of evaluation unchanged and outputs the item at the current position.

#### Print$(n)$

The instruction

$$\text{Print}(n)$$

outputs item $n$ without changing the point of evaluation.

```
def print_item(n, position):
    return str(n), position
```

For example,

```
Print(2)
Print(2)
Print(3)
```

produces

$$223$$

independently of the initial point of evaluation.

### Concatenation

Programs can be composed sequentially by concatenation. If $P$ and $Q$ are programs, their concatenation

$$P; Q$$

means: first execute $P$, and then execute $Q$ starting from the point of evaluation at which $P$ ended. The output of $P; Q$ is the output of $P$ followed by the output of $Q$.

More generally, for

$$P = P_1; P_2; \dots; P_r,$$

the subprograms are executed from left to right. Both components of the machine state are threaded through the concatenation: the output produced by each $P_i$ is appended to the output accumulated so far, and the final point of evaluation of $P_i$ becomes the initial point of evaluation of $P_{i+1}$.

```
def concatenate(programs, position, base):
    output = ""

    for P in programs:
        block, position = execute(P, position, base)
        output += block

    return output, position
```

For example, with $b = 6$, the concatenation

```
Move&Print(+1)
Move&Print(+2)
```

executed from $x = 0$ first produces $1$ and leaves the point of evaluation at $1$. The second instruction therefore starts from $1$, moves to $3$, and produces $3$. The complete output is

$$13.$$

Concatenation is the basic mechanism by which instructions and subprograms are combined into larger programs.

### Repeat

The construct

$$\text{Repeat}(k, P)$$

executes program $P$ exactly $k$ consecutive times. The point of evaluation is carried from one execution of $P$ to the next.

```
def repeat(k, P, position, base):
    output = ""

    for _ in range(k):
        block, position = execute(P, position, base)
        output += block

    return output, position
```

For example, with $b = 6$,

```
Repeat 4 times:
    Move&Print(+1)
```

executed from $x = 0$ produces

$$1234,$$

whereas from $x = 4$ it produces

$$5012.$$

Repeats may be nested. For example,

```
Repeat 3 times:
    Move&Print(+1)
    Repeat 2 times:
        Move&Print(0)
```

executed from $x = 0$ produces

$$111222333.$$

### Repeat with application

The second repetition construct has the form

$$\text{RepeatApply}\big(k, P, \text{Move\&Print}(m)\big).$$

It first executes $P$ once from the current point of evaluation. Suppose that this first execution produces the block

$$a_1 a_2 \cdots a_r.$$

This block forms the first part of the output.

The remaining copies are obtained in a different way from an ordinary $\text{Repeat}$. The program $P$ is not executed again. Instead, the instruction $\text{Move\&Print}(m)$ is applied separately to each symbol of the block that has just been produced. Thus, to construct the next block, $a_i$ is taken as a point of evaluation, $\text{Move\&Print}(m)$ is executed from $a_i$, and the resulting output symbol becomes the $i$th symbol of the new block. Once the whole new block has been obtained, the same operation is applied to its symbols to generate the following block. This process is repeated until $k$ blocks have been produced.

The final point of evaluation is the last symbol in the resulting output.

The following Python-like pseudocode summarizes this behavior:

```
def repeat_apply(k, P, application, position, base):
    block, _ = execute(P, position, base)
    output = block

    for _ in range(k - 1):
        next_block = ""

        for symbol in block:
            printed, _ = execute(
                (application,),
                int(symbol),
                base
            )
            next_block += printed

        block = next_block
        output += block

    return output, int(output[-1])
```

For example,

```
Repeat 3 times with application of Move&Print(+1):
    Repeat 3 times:
        Move&Print(+1)
```

executed from $x = 0$ first produces

$$123.$$

Applying $\mathrm{Move\&Print}(+1)$ independently to the three symbols produces

$$234,$$

and applying the same instruction to the symbols of this new block produces

$$345.$$

Hence the complete output is

$$123234345.$$

### Back to start

The paper uses *Back to start* for the special case

$$\mathrm{RepeatApply}\big(k, P, \mathrm{Move\&Print}(0)\big).$$

Because $\mathrm{Move\&Print}(0)$ leaves each point unchanged, each block following the first one is identical to the preceding block. Thus the same block is reproduced at every repetition.

For example,

```
Repeat 4 times, Back to start:
    Repeat 3 times:
        Move&Print(+1)
```

denotes

```
Repeat 4 times with application of Move&Print(0):
    Repeat 3 times:
        Move&Print(+1)
```

and, from $x = 0$, produces

$$123123123123.$$

*Back to start* is therefore an abbreviated name for this special case and does not introduce an additional primitive or an additional weight.

### Program size and LoT-complexity

Each primitive has an associated weight. Let $L(P)$ denote the size of program $P$. For the elementary instructions,

$$L(\mathrm{Move\&Print}(n)) = W_{\mathrm{Move\&Print}}(n),$$

and

$$L(\mathrm{Print}(n)) = W_{\mathrm{Print}}(n).$$

If

$$P = P_1; \dots ; P_r,$$

then program size is additive:

$$L(P) = \sum_{i=1}^{r} L\,(P_i).$$

For repetition,

$$L(\mathrm{Repeat}(k, P)) = W_{\mathrm{Repeat}}(k) + L(P).$$

The repeat-with-application construct has the additional cost of the instruction applied between blocks:

$$L\left(\mathrm{RepeatApply}(k, P, \mathrm{Move\&Print}(m))\right)$$
$$= W_{\mathrm{Repeat}}(k) + W_{\mathrm{Move\&Print}}(m) + L(P).$$

Consequently, *Back to start* has size

$$W_{\mathrm{Repeat}}(k) + W_{\mathrm{Move\&Print}}(0) + L(P).$$

Let $\mathrm{out}(P, x)$ be the output of program $P$ when its initial point of evaluation is $x$. The LoT-complexity of a sequence $s$ is the minimum weighted program size among all programs and all possible initial points of evaluation that generate $s$:

$$C_{\mathrm{LoT}}(s) = \min_{\substack{P,\ x \in \mathcal{A}_b \\ \mathrm{out}(P,x)=s}} L(P).$$

The minimal program therefore depends on the primitive weights: changing the weights changes the relative sizes of candidate programs and may change which description is minimal.

### Primitive weights in Experiment 2

Experiment 2 of the paper compares a fixed-weight baseline with two parametrizations in which the primitive weights depend on their arguments.

For the baseline LoT model,

$$W_{\mathrm{Repeat}}(n) = 1,$$

and

$$W_{\mathrm{Move\&Print}}(n) = 2.$$

For Model 1,

$$W_{\text{Repeat}}(n) = 1 + \beta\log_2(n),$$

and

$$W_{\text{Move\&Print}}(n) = 1 + \alpha|n|.$$

For Model 2,

$$W_{\text{Repeat}}(n) = \beta\log_2(n) + \gamma,$$

while

$$W_{\text{Move\&Print}}(n) = 1 + \alpha|n|.$$

Here $n$ in $W_{\text{Repeat}}(n)$ is the number of repetitions, whereas $n$ in $W_{\text{Move\&Print}}(n)$ is the signed ordinal displacement. The absolute value $|n|$ makes the movement cost depend on ordinal distance but not on direction.

The parameters are fitted as described in Experiment 2. For every choice of the weights, LoT-complexity is obtained by searching for the minimum-size program under those weights. The weight search and the minimal-program search are therefore linked: a change in primitive weights can change both the numerical complexity of a sequence and the program that achieves it.

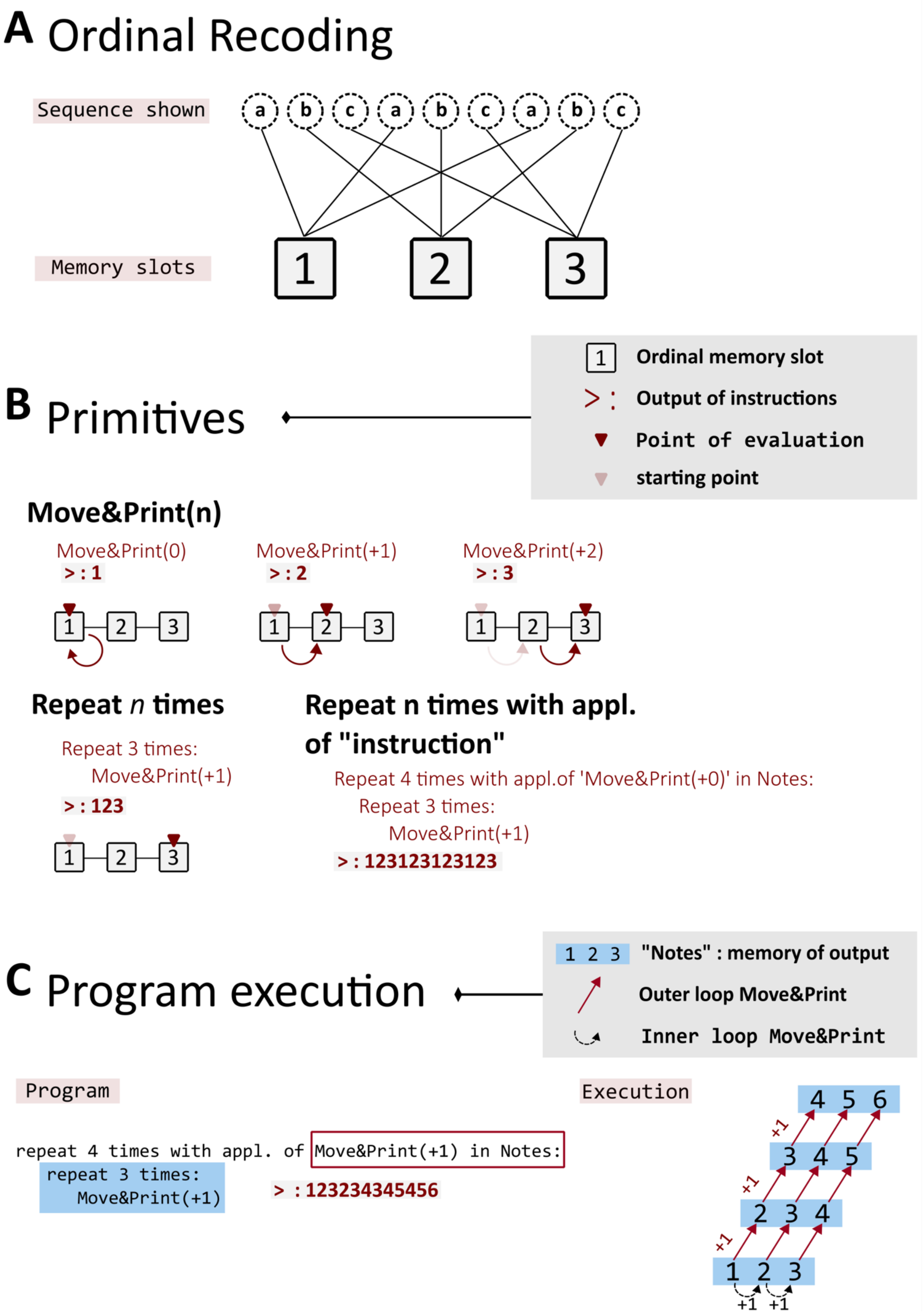


**Fig A. Implementing the Language of Thought (LoT) framework.**
**(A)** Ordinal recoding. The visuo-spatial sequence is recoded into ordinal memory slots according to each item's order of first appearance. Because spatial positions are randomly assigned on every trial, distinct spatial sequences (here, abcabcabc) that share the same order-

based structure map onto the same ordinal expression (here, 123123123). **(B)** Primitives. The LoT composes programs from a small set of primitives operating over ordinal memory slots. "Move&Print(n)" advances the point of evaluation by n slots and outputs the element at that position; "Move&Print(0)" outputs the current slot without advancing. "Repeat n times" applies an instruction a fixed number of times, and can itself take an instruction applied at each iteration in the version "Repeat n times with appl. of 'instruction'", allowing nested loops with variation. By convention, the point of evaluation is initialized at position 0. **(C)** Program execution. A nested program (left) and its step-by-step execution (right). The inner loop (Repeat 3 times: Move&Print(+1)) generates a three-element chunk, which is stored in Notes, the memory of the output; the outer loop then re-executes this chunk four times, applying Move&Print(+1) to the stored chunk at each iteration so that successive chunks are shifted by one ordinal position, yielding 123234345456.

**Fig B. Distribution of sequence lengths of error trials obtained for the sequences of Experiment 1.**

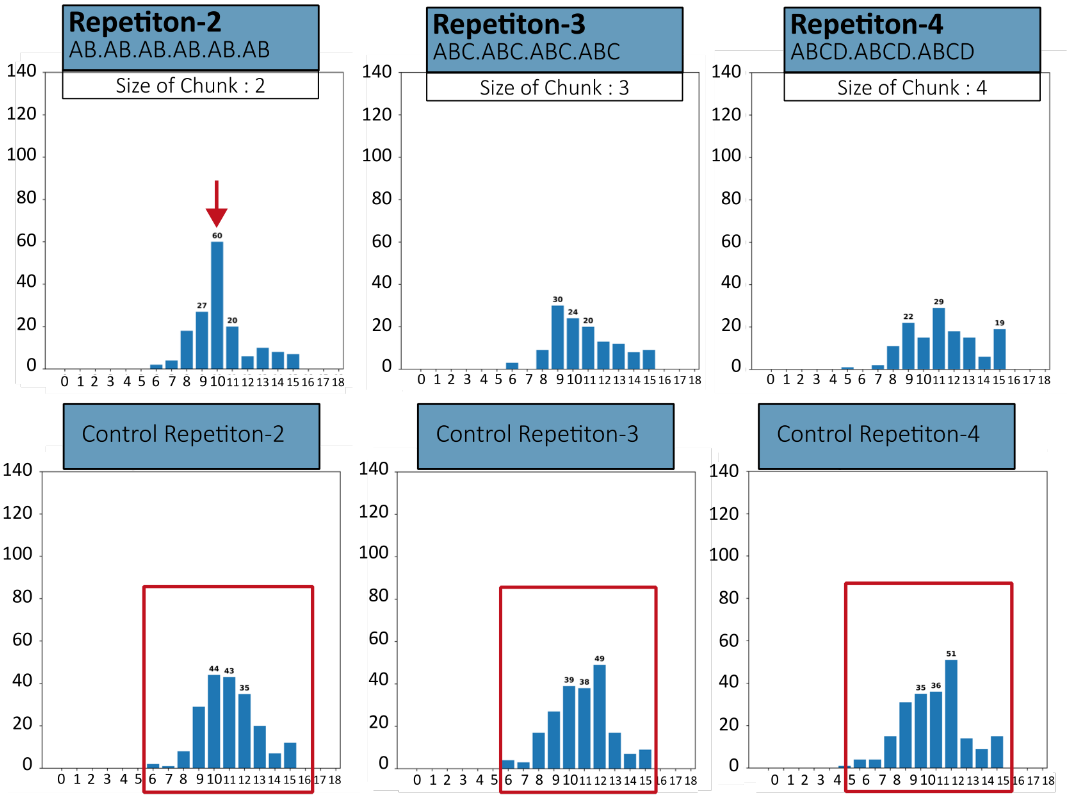


Arrows indicate peaks in response length, suggesting that participants tend to forget groups of items (chunks) rather than single items. Grouped forgetting appears in sequences with four or more chunks but is absent in control sequences, suggesting that participants hold a noisy representation of the number of repetitions ($n$) of a program ($P$).

**Fig C. Inter-click timings and cumulative accuracy for Experiment 2 full sequence.**

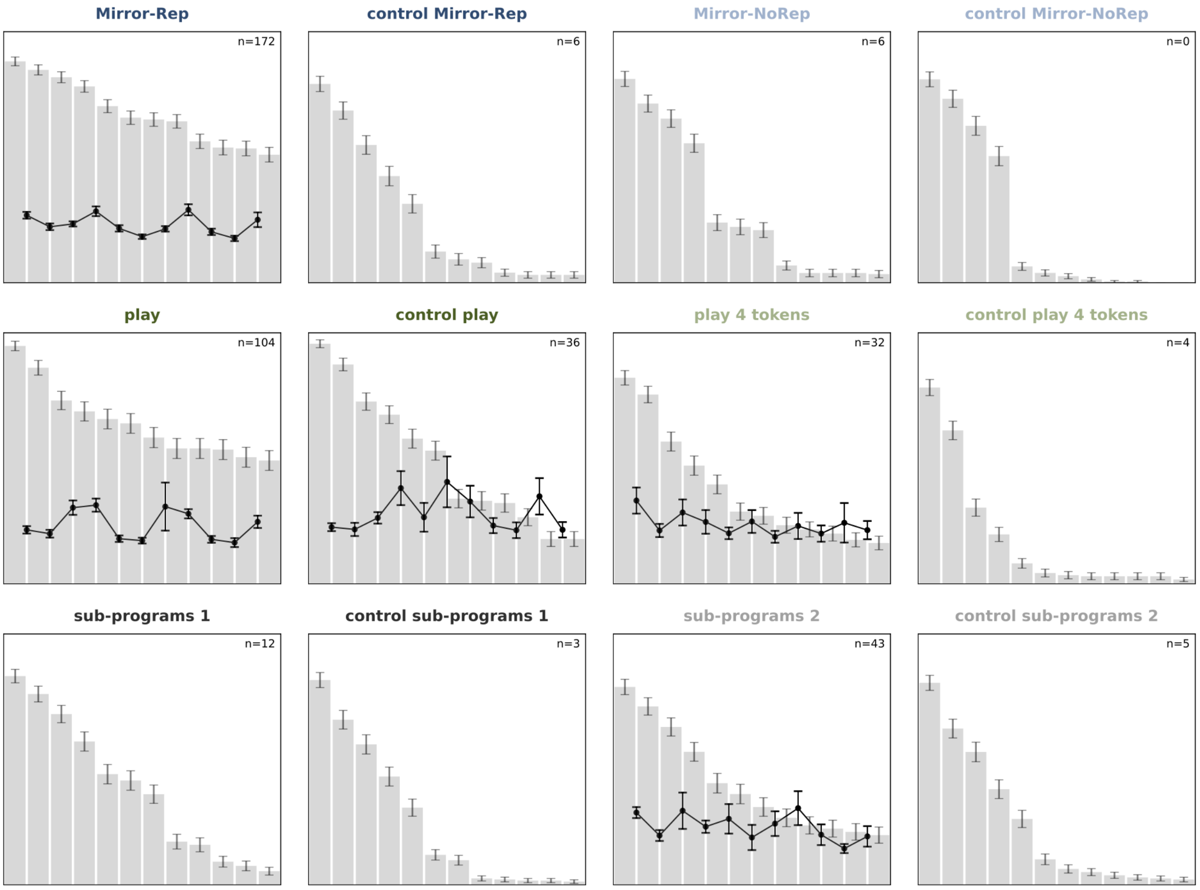


The figure overlays (1) cumulative accuracy (background histograms) and (2) inter-click intervals computed on accurately reproduced sequences. Error bars represent the standard error of the mean. The number of correct responses for each sequence (Experiment 2 only) is displayed in the top-right corner of each panel. Only sequences with n $\geq$ 20 correct trials were considered for inter-click intervals plotting. Drops in cumulative accuracy and peaks in inter-click durations mark chunk boundaries (Kennerley et al., 2004).

**Table A. LoT description of the exploratory sequences MirrorRep, NamedSubprogram-2 and Print-4.**

| ***Sequence Name*** | **MirrorRep** | **NamedSubprogram-2** | **Print-4** |
|---|---|---|---|
| ***LoT proposed minimal program*** | ***repeat 4 times:***<br>***move +1 and play***<br>***move 0 and play***<br>***repeat 3 times:***<br>***move -1 and play***<br>***move 0 and play***<br>***repeat 3 times:*** | ***move 0 and play***<br>***repeat 2 times with application***<br>***of 'move -1 and play' in notes:***<br>***repeat 3 times:***<br>***move +1 and play*** | ***repeat 2 times with application***<br>***of 'move 0' in notes:***<br>***move 0 and play***<br>***move +1 and play***<br>***move -1 and play***<br>***move +2 and play*** |

| | | | |
|---|---|---|---|
| | *move +1 and play* | *repeat 2 times:*<br>*move +2 and play*<br>*repeat 2 times:*<br>*move +1 and play*<br>*move +3 and play* | *move -2 and play*<br>*move +3 and play* |
| ***Sequence expression*** | **ABCDDCBAABCD** | **ABCDABCEABCF** | **ABACADABACAD** |

**Table B. Mean error rate per sequence (Experiment 1)**

| Condition | n | Mean Error Rate | SEM |
|---|---|---|---|
| Rep-2 | 82 | 75.61 | 4.00 |
| CRep-2 | 82 | 95.73 | 2.16 |
| Rep-3 | 82 | 60.37 | 4.40 |
| CRep-3 | 82 | 97.56 | 1.48 |
| Rep-4 | 82 | 64.02 | 4.27 |
| CRep-4 | 82 | 99.39 | 0.61 |
| Rep-Nested | 82 | 53.66 | 4.65 |
| Rep-Global | 82 | 91.46 | 2.72 |
| Rep-Local | 82 | 70.12 | 4.24 |
| Overall Average | 82 | 78.66 | 1.31 |

**Table C. Mean error rate paired-comparisons (Experiment 1)**

| Comparison (A vs. B) | Non-zero Pairs | Wilcoxon W | p-value | Effect Size (r) |
|---|---|---|---|---|
| Rep-2 vs. CRep-2 | 27 | 10.00 | p<.001 | 0.87 (Large) |
| Rep-3 vs. CRep-3 | 45 | 0.00 | p<.001 | 0.91 (Large) |
| Rep-4 vs. CRep-4 | 43 | 0.00 | p<.001 | 0.91 (Large) |
| Rep-Nested vs. Rep-3 | 40 | 310.00 | p=.142 | 0.23 (Small) |
| Rep-Nested vs. CRep-3 | 48 | 0.00 | p<.001 | 0.90 (Large) |
| Rep-Nested vs. Rep-Global | 48 | 43.50 | p<.001 | 0.83 (Large) |
| Rep-Nested vs. Rep-Local | 45 | 265.50 | p=.003 | 0.45 (Medium) |
| Rep-Global vs. CRep-3 | 9 | 0.00 | p=.004 | 0.96 (Large) |

| | | | | |
|---|---|---|---|---|
| **Rep-Local vs. CRep-3** | **34** | **0.00** | **p<.001** | **0.91 (Large)** |
| **Rep-Global vs. Rep-Local** | **28** | **10.00** | **p<.001** | **0.87 (Large)** |

**Table D. Mean error rate group comparison (Experiment 1)**

| **Group** | **Mean (%)** | **SEM** | **Wilcoxon W** | **p-value** | **Effect Size (r)** |
|---|---|---|---|---|---|
| **Structured** | **66.67** | **3.11** | **0.00** | **p<.001** | **0.88 (Large)** |
| **Control** | **97.56** | **1.23** | | | |

**Table E. Mean Edit distance per sequence (Experiment 1)**

| Sequence Name | N | Mean DL | SEM |
|---|---|---|---|
| Rep-2 | 82 | 1.8000 | 0.1317 |
| CRep-2 | 82 | 3.5900 | 0.1386 |
| Rep-3 | 82 | 1.6600 | 0.1557 |
| CRep-3 | 82 | 4.4100 | 0.1269 |
| Rep-4 | 82 | 2.4000 | 0.2082 |
| CRep-4 | 82 | 5.6500 | 0.1291 |
| Rep-Nested | 82 | 1.5900 | 0.1705 |
| Rep-Global | 82 | 3.7200 | 0.1494 |
| **Rep-Local** | 82 | 2.9600 | 0.2258 |

**Table F. Mean Edit distance paired comparisons (Experiment 1)**

| **Comparison Pair** | **n** | **W** | **p-value** | **r** | **Effect Size** |
|---|---|---|---|---|---|
| **Rep-2 vs. CRep-2** | 82 | 29.00 | 0.0000 | 0.8530 | large *** |
| **Rep-3 vs. CRep-3** | 82 | 8.50 | 0.0000 | 0.8658 | large *** |
| **Rep-4 vs. CRep-4** | 82 | 21.00 | 0.0000 | 0.8587 | large *** |
| **Rep-Nested vs. Rep-3** | 82 | 895.00 | 0.5652 | 0.0730 | small ns |
| **Rep-Nested vs. CRep-3** | 82 | 14.00 | 0.0000 | 0.8624 | large *** |
| **Rep-Nested vs. Rep-Global** | 82 | 101.50 | 0.0000 | 0.8115 | large *** |
| **Rep-Nested vs. Rep-Local** | 82 | 452.00 | 0.0000 | 0.5874 | large *** |
| **Rep-Global vs. CRep-3** | 82 | 420.50 | 0.0000 | 0.5526 | large *** |
| **Rep-Local vs. CRep-3** | 82 | 432.50 | 0.0000 | 0.6065 | large *** |
| **Rep-Global vs. Rep-Local** | 82 | 719.00 | 0.0021 | 0.3677 | medium ** |

**Table G. Mean Edit distance group comparisons (Experiment 1)**

| Group | n | Mean DL | SEM | W | p-value | r | Effect Size |
|---|---|---|---|---|---|---|---|
| **Structured** | 82 | 1.9533 | 0.1229 | 0.00 | 0.0000 | 0.8688 | large *** |
| **Control** | 82 | 4.5508 | 0.1029 | — | — | — | — |

**Table H. Mean Total response time per sequence (Experiment 1)**

| Sequence Name | N | Mean (ms) | SEM (ms) |
|---|---|---|---|
| **Rep-2** | 82 | 5728.77 | 160.66 |
| **CRep-2** | 82 | 6526.69 | 157.21 |
| **Rep-3** | 82 | 6154.74 | 151.04 |
| **CRep-3** | 82 | 6512.85 | 180.14 |
| **Rep-4** | 82 | 6557.09 | 171.79 |
| **CRep-4** | 82 | 6441.40 | 190.84 |
| **Rep-Nested** | 82 | 6232.38 | 147.34 |
| **Rep-Global** | 82 | 6638.32 | 190.86 |
| **Rep-Local** | 82 | 6672.71 | 178.58 |
| **Overall (all sequences)** | — | 6384.99 | 57.80 |

**Table I. Mean Total response time paired comparisons (Experiment 1)**

| Comparison Pair | n | W | p-value | r | Effect Size |
|---|---|---|---|---|---|
| **Rep-2 vs. CRep-2** | 82 | 661.50 | 0.0000 | 0.5309 | large *** |
| **Rep-3 vs. CRep-3** | 82 | 1142.00 | 0.0097 | 0.2856 | small ** |
| **Rep-4 vs. CRep-4** | 82 | 1487.50 | 0.3225 | 0.1093 | small ns |
| **Rep-Nested vs. Rep-3** | 82 | 1608.00 | 0.6656 | 0.0477 | small ns |
| **Rep-Nested vs. CRep-3** | 82 | 1313.50 | 0.0729 | 0.1981 | small ns |
| **Rep-Nested vs. Rep-Global** | 82 | 1243.50 | 0.0342 | 0.2338 | small * |
| **Rep-Nested vs. Rep-Local** | 82 | 1002.00 | 0.0012 | 0.3571 | medium ** |
| **Rep-Global vs. ctrl Rep-3** | 82 | 1560.00 | 0.5130 | 0.0722 | small ns |
| **Rep-Local vs. ctrl Rep-3** | 82 | 1520.00 | 0.4014 | 0.0927 | small ns |

| | | | | | |
|---|---|---|---|---|---|
| **Rep-Global vs. Rep-Local** | 82 | 1625.00 | 0.7236 | 0.0391 | small ns |

**Table J. Mean Total response time group comparisons (Experiment 1)**

| Group | n | Mean (ms) | SEM | W | p-value | r | Effect Size |
|---|---|---|---|---|---|---|---|
| **Structured** | 82 | 6146.87 | 141.10 | 1047.00 | 0.0025 | 0.3341 | medium ** |
| **Control** | 82 | 6493.65 | 150.17 | — | — | — | — |

**Table K. Response Length – Errors Only (Experiment 1)**

| Sequence | N Trial | N PP | Mean | SD | Mode | Variance |
|---|---|---|---|---|---|---|
| **Rep-2** | 124 | 71 | 10.43 | 2.00 | 10 | 3.9544 |
| **CRep-2** | 157 | 79 | 11.03 | 1.81 | 11 | 3.2410 |
| **Rep-3** | 99 | 63 | 10.81 | 2.10 | 9 | 4.3773 |
| **CRep-3** | 160 | 81 | 10.82 | 1.97 | 12 | 3.8569 |
| **Rep-4** | 105 | 66 | 11.35 | 2.33 | 11 | 5.3711 |
| **CRep-4** | 163 | 82 | 10.80 | 2.11 | 12 | 4.4154 |
| **Rep-Nested** | 88 | 56 | 11.40 | 2.34 | 10 | 5.4214 |
| **Rep-Global** | 150 | 78 | 10.83 | 1.99 | 12 | 3.9522 |
| **Rep-Local** | 115 | 68 | 11.74 | 2.06 | 12 | 4.1928 |

SD = Standard Deviation
Kurt = Kurtosis
N Trial = Number of error trials available for analysis (after removing correct and length ≥15)
N PP = Number of unique participants contributing a response for length analysis.

**Table L. Response Length – Centrality**
One-sample t-test vs μ=12 and Cohen's d Unit of analysis: per-participant mean response length.

| Sequence | N_pp | Mean_pp | SD_pp | t | p | Cohen d |
|---|---|---|---|---|---|---|
| **Rep-2** | 71 | 10.521 | 1.831 | -6.805 | 0.0000* | -0.808 |
| **CRep-2** | 79 | 11.051 | 1.499 | -5.629 | 0.0000* | -0.633 |
| **Rep-3** | 63 | 10.984 | 1.974 | -4.086 | 0.0001* | -0.515 |
| **CRep-3** | 81 | 10.840 | 1.631 | -6.402 | 0.0000* | -0.711 |
| **Rep-4** | 66 | 11.447 | 2.083 | -2.157 | 0.0347* | -0.265 |
| **CRep-4** | 82 | 10.811 | 1.739 | -6.193 | 0.0000* | -0.684 |
| **Rep-Nested** | 56 | 11.518 | 2.172 | -1.661 | 0.1024 | -0.222 |
| **Rep-Global** | 78 | 10.865 | 1.699 | -5.897 | 0.0000* | -0.668 |
| **Rep-Local** | 68 | 11.772 | 1.888 | -0.996 | 0.3229 | -0.121 |

**Fig D. Distribution of all responses for experiment 1.**

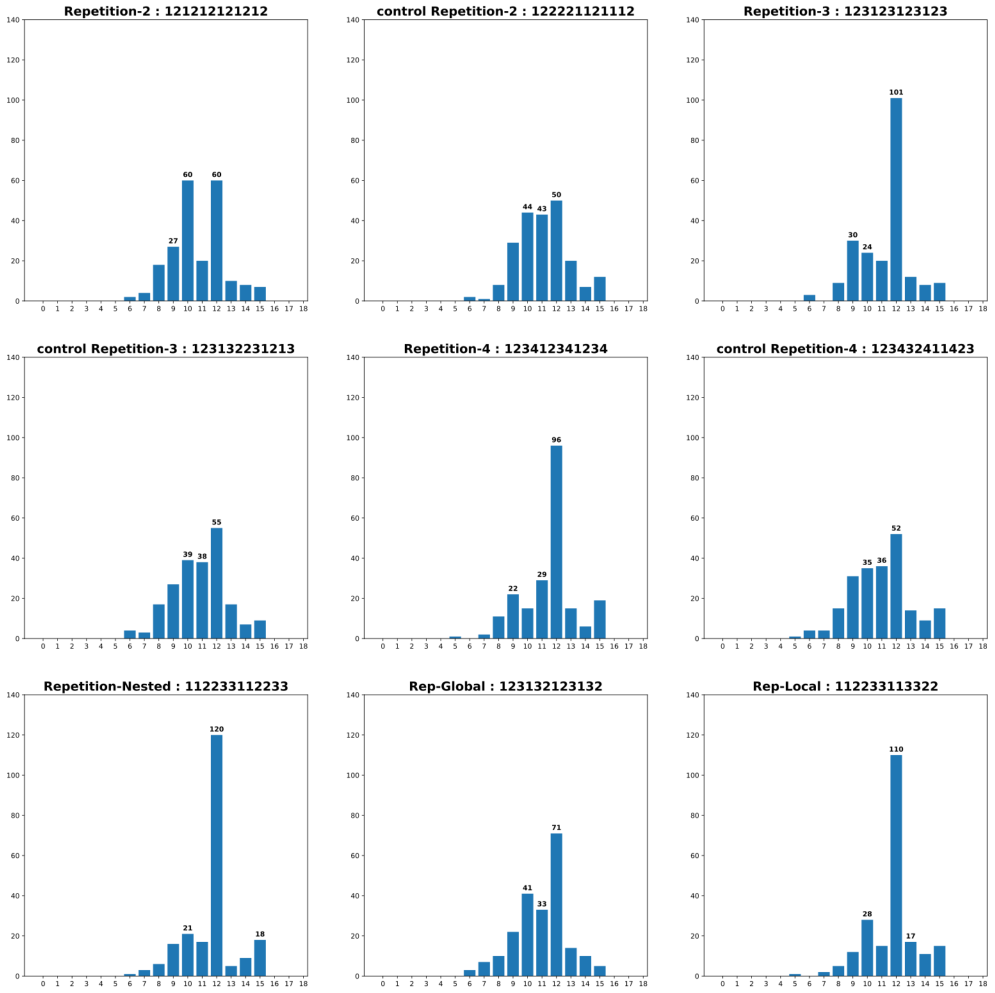
Repetition-2 : 121212121212
27
60
60
control Repetition-2 : 122221121112
44
43
50
Repetition-3 : 123123123123
30
24
101
control Repetition-3 : 123132231213
39
38
55
Repetition-4 : 123412341234
22
29
96
control Repetition-4 : 123432411423
35
36
52
Repetition-Nested : 112233112233
21
120
18
Rep-Global : 123132123132
41
33
71
Rep-Local : 112233113322
28
110
17

### Table M. Retroactive interference measures (Experiment 1)

We computed the error rate of only the first chunks in the participant responses. The length of first chunk was determined according to our defined LoT description of the LoT structured sequence and extended to its control(s) for comparability. Even though the LoT description of controls differed, LoT structured sequences and their controls start exactly in the same way. Which allows for comparisons of the effect of structure on a group of first items. Exception for Nested Repetition and control NoLocal. Statistical test: Wilcoxon Signed-Ranked Test

| Sequence Name | n-first items considered | Structured - Mean chunk error rate | SEM | Control - Mean chunk error rate | SEM | Difference | Test-stat | P-Value | Effect size (r = Z / √N) |
|---|---|---|---|---|---|---|---|---|---|
| Rep-2 | AB | 18.95 | 2.9 | 34.21 | 3.85 | 15.26 | 246.0 | <0.001 | 0.474 |
| Rep-3 | ABC | 26.22 | 3.80 | 57.93 | 3.93 | 31.71 | 312.0 | <0.001 | 0.641 |
| Rep-4 | ABCD | 40.24 | 4.46 | 66.46 | 3.98 | 26.22 | 136.0 | <0.001 | 0.685 |
| Rep-Nested / Rep-Local | AABBCC | 28.68 | 3.97 | 41.32 | 3.95 | 12.63 | 167.0 | <0.0001 | 0.725 |

### Table N. Retroactive interference measures (Experiment 2)

| Sequence Name | n-first items considered | Structured - Mean chunk error rate | SEM | Control - Mean chunk error rate | SEM | Difference | Test-stat | P-Value | Effect size (r = Z / √N) |
|---|---|---|---|---|---|---|---|---|---|
| Rep-2 | AB | 13.96 | 2.79 | 26.99 | 3.37 | 13.03 | 359.0 | 0.00469 | 0.38 |
| Rep-3 | ABC | 13.84 | 2.46 | 42.54 | 3.84 | 28.7 | 384.0 | 5.45e-08 | 0.615 |
| Rep-4 | ABCD | 34.78 | 3.65 | 54.5 | 3.69 | 19.72 | 825.0 | 0.00032 | 0.391 |
| Rep-Nested | AABBCC | 18.38 | 3.29 | 34.48 | 3.52 | 16.11 | 333.5 | <0.0001 | 0.636 |
| Print 4 Tokens | ABAC | 52.92 | 3.83 | 80.34 | 2.83 | 27.43 | 514.0 | 3.17e-07 | 0.564 |
| Sub-Programs 1 | ABC | 31.93 | 3.45 | 44.02 | 3.83 | 12.08 | 1010.5 | 0.02281 | 0.253 |
| Sub-Programs 2 | ABCD | 46.98 | 3.72 | 61.86 | 3.73 | 14.88 | 925.0 | 0.00595 | 0.305 |
| Print | AAAB | 31.3 | 3.81 | 32.61 | 3.53 | 1.3 | 1256.0 | 0.89576 | 0.015 |
| Mirror-Rep | ABCD | 21.79 | 2.3 | 57.52 | 3.87 | 35.73 | 593.5 | 1.35e-09 | 0.627 |
| Mirror-NoRep | ABCD | 44.44 | 3.61 | 49.57 | 3.83 | 5.12 | 1103.5 | 0.29871 | 0.119 |
